# Massively parallel assessment of designed protein solution properties using mass spectrometry and peptide barcoding

**DOI:** 10.1101/2025.02.24.639402

**Authors:** David Feldman, Jeremiah N. Sims, Xinting Li, Richard Johnson, Stacey Gerben, David E. Kim, Christian Richardson, Brian Koepnick, Helen Eisenach, Derrick R. Hicks, Erin C. Yang, Basile I. M. Wicky, Lukas F. Milles, Asim K. Bera, Alex Kang, Evans Brackenbrough, Emily Joyce, Banumathi Sankaran, Joshua M. Lubner, Inna Goreshnik, Dionne Vafeados, Aza Allen, Lance Stewart, Michael J. MacCoss, David Baker

## Abstract

Library screening and selection methods can determine the binding activities of individual members of large protein libraries given a physical link between protein and nucleotide sequence, which enables identification of functional molecules by DNA sequencing. However, the solution properties of individual protein molecules cannot be probed using such approaches because they are completely altered by DNA attachment. Mass spectrometry enables parallel evaluation of protein properties amenable to physical fractionation such as solubility and oligomeric state, but current approaches are limited to libraries of 1,000 or fewer proteins. Here, we improved mass spectrometry barcoding by co-synthesizing proteins with barcodes optimized to be highly multiplexable and minimally perturbative, scaling to libraries of >5,000 proteins. We use these barcodes together with mass spectrometry to assay the solution behavior of libraries of *de novo*-designed monomeric scaffolds, oligomers, binding proteins and nanocages, rapidly identifying design failure modes and successes.

## Main

Rapid improvements in computational methods have enabled the design of proteins with increasingly sophisticated structure and function (Watson et al. 2023; Dauparas et al. 2022; Baek et al. 2021). Nevertheless, for many tasks the success rate remains low, and the deficiencies in computational models are unclear. Library screens address this gap via efficient pooled testing of thousands of designs, identifying candidates for further development and providing feedback to improve the design process. A wide variety of enrichment methods have been applied to protein libraries, including display methods to measure binding affinity (Boder and Wittrup 1997) and protease stability (Tsuboyama et al. 2023), fluorescent substrate turnover for enzymatic activity (Harris et al. 2000), and split GFP or enzymatic reporters as proxies for solubility (Cabantous and Waldo 2006; Zutz et al. 2021). However, these methods rely on physical linkage between the protein and the DNA sequence which encodes it, and this requirement has precluded directly screening proteins in solution for key biophysical properties such as oligomeric state, solubility, expression yield, and functional delivery (Egloff et al. 2019; Rhym et al. 2023). Mass spectrometry, by contrast, enables parallel evaluation of protein properties such as aggregation propensity and resistance to chemical or thermal denaturation in solution. Physical fractionation methods such as size exclusion chromatography or bead-based pulldown of substrate-bound protein can be performed on a sample containing many proteins of interest, and the identities of the proteins in each fraction determined by mass spectrometry. However, direct application of shotgun proteomics to large libraries of designed proteins is frequently limited by high sequence similarity among designs. To circumvent sequence similarity, mass spectrometry multiplexing via peptide barcodes was recently developed to evaluate nanobody solution binding, monomericity, and expression (Egloff et al. 2019). However, this method used stochastic linking of peptide barcodes with randomized sequences to designs, and was thus limited to a pre-enriched pool of 1000 nanobodies attached to ∼12,000 total barcodes.

We reasoned that an approach combining mass spectrometry with peptide barcodes could provide a powerful way of assessing the properties of thousands of designed proteins in solution. We set out to optimize mass spectrometry barcoding for measuring the properties of diverse *de novo*-designed proteins, including monomeric and oligomeric scaffolds, minibinders, and nanocages. Since the previous approach used a shotgun cloning strategy to stochastically assign barcodes to a small nanobody library, there was an inherent need to use next generation sequencing (NGS) to identify unique barcode assignments to designs. While the previous approach proved effective, we reasoned that pre-assignment of barcodes at the DNA oligo level would afford greater overall throughput than a shotgun cloning approach, because pre-assignment would mitigate non-unique pairings of barcode and designed protein. Additionally, the previous approach suffered from barcode dropout at the level of mass spectrometry, likely due to barcode-specific differences in ionization efficiency. Thus, we aimed to identify a set of barcodes that would ionize reliably, thereby further increasing the fidelity of barcode identification, and subsequently, improving throughput. We began by seeking to design peptide barcodes *in silico* that are (1) co-synthesized with designs on an oligonucleotide array; (2) easily purified for mass spectrometry (Figure 1a); (3) minimally perturbative to the attached designs; and (4) efficiently separated and quantified by high resolution orbitrap liquid chromatography-coupled tandem mass spectrometry (LC-MS/MS) (Figure 1b).

**Figure 1:**
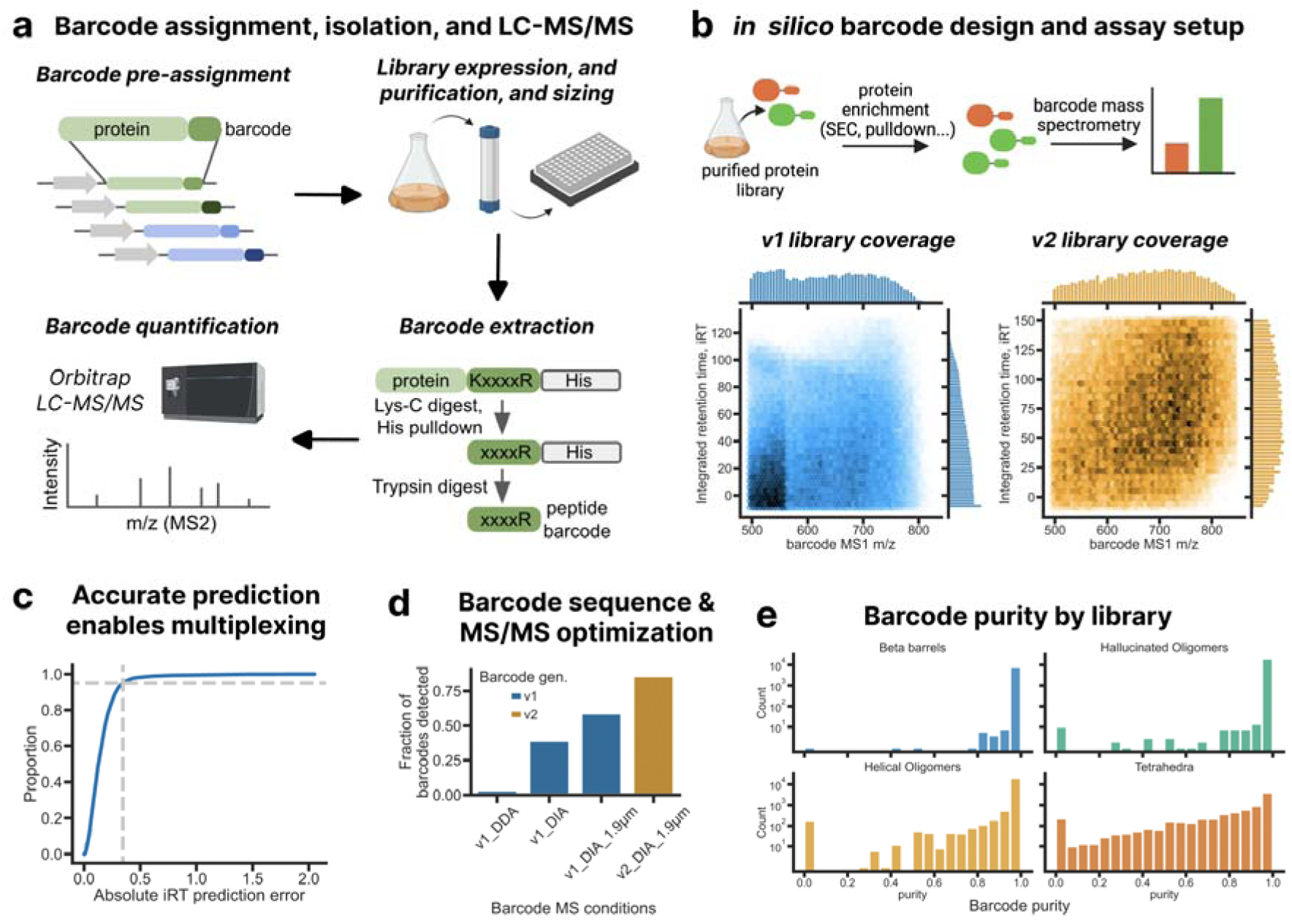
Barcode design and characterization. **(a)** Schematic of mass spectrometry barcoding workflow. Orthogonal barcode sets are designed *in silico* based on predicted LC-MS/MS performance and biophysical properties. Designed protein-barcode pairs are synthesized as an oligo pool for proteins up to 70 aa (154 aa with oligo assembly). The protein library is then expressed and fractionated as a single sample. Barcodes are purified from each fraction and measured by orbitrap LC-MS/MS. **(b)** Protein enrichments are estimated from normalized barcode intensity, which requires efficient discrimination of barcodes. LC-MS/MS permits separation of barcodes by hydrophobicity (which influences iRT) and by parent ion mass-to-charge ratios (m/z). Barcodes were restricted to 450-850 m/z to remain within the optimal m/z range for DDA/DIA Orbitrap while spanning the space of iRT. **(c)** Accuracy of barcode elution from the LC column was measured empirically by comparing the predicted iRT (via Prosit) and the experimentally measured iRT (via DIA-NN) for the v2 barcoding set tagged to muGFP. A difference of < 0.5 iRT between actual and predicted iRT was observed for 95% of barcodes (grey dotted lines), demonstrating high accuracy for *in silico* prediction of barcode LC retention. **(d)** An improved barcode detection rate was achieved across benchmarking muGFP barcoding experiments by (1) switching from DDA to DIA, (2) employing columns packed with finer silica (5 µm → 1.9 µm), and (3) restricting the amino acid composition of barcodes. **(e)** The frequency that a barcode is correctly mapped to a design (barcode purity) for different libraries. For the single oligo libraries, 99% of barcodes exclusively mapped to the intended design, and ∼80% for 2-oligo designs (requiring pooled PCR homologous assembly).

To meet criteria (1) and (2), we adapted pET-28, a T7 expression vector, for library cloning of 300-nt oligos containing barcodes fused to protein designs of up to 74 amino acids, or 154 amino acids if using oligo assembly. Proteins expressed from this vector contain a barcode flanked by a designed protein and either an N- or C-terminal His-tag. Arginine and lysine residues were restricted to the barcode boundaries to enable facile isolation of barcodes from the protein of interest and tags by sequential protease digest by LysC digest, His-tag purification, and then trypsin digest. Barcodes were limited to 8 to 13 amino acids in length and predicted to generate doubly-charged precursors by electrospray ionization, in order to optimally cover the mass-to-charge range of high resolution orbitrap mass spectrometry. To meet criterion (3), the amino acid content of the barcodes was limited to avoid bulky hydrophobic residues likely to disrupt folding, as well as residues that affect net charge in electrospray ionization, residues likely to undergo chemical modification, and residues that interfere with tryptic cleavage (Methods). Finally, to meet criterion (4), LC indexed retention time (iRT, a standardized measurement for predicting elution, (Escher et al. 2012)) and MS/MS fragmentation spectra were predicted for candidate barcodes using Prosit, and a first-generation set of up to 100,000 barcodes (v1 barcodes) was defined based on separability at the expected m/z and LC resolution.

As an initial test, a subset of 5,000 first-generation barcodes was synthesized and appended to Foldit1, a highly stable and soluble protein characterized previously (Koepnick et al. 2019). The barcodes were expressed in subpools and mixed to generate a standard curve for ratiometric quantification. After expression, barcodes were purified directly from *E. coli* lysate and analyzed by data-dependent acquisition (DDA) on an Orbitrap mass spectrometer (Methods), and barcode intensities were quantified by integrated MS1 area (Methods).

Application of the v1 barcodes to the larger design libraries in this study revealed that fewer barcoded designs were identified via mass spectrometry than by DNA sequencing (∼30% detection rate across libraries with >10,000 barcodes), similar to the barcode design detection rate reported in Egloff, 2019. We suspected that this detection failure might be due to both unstable designs and suboptimal barcode sequence content for our mass spectrometry protocol; to isolate the latter scenario, we barcoded muGFP, a hyperstable monomeric variant of green fluorescent protein. We sought to improve the barcode detection rate by optimizing the sensitivity of our mass spectrometry protocol and devising a second-generation barcode set (v2 barcodes) based on the experimental data. Stochastic detection and quantification is a known limitation of data-dependent acquisition (DDA) (Barkovits et al. 2020), which relies on ion abundance in MS1 scans to trigger MS2 scans for peptide identification. This focus on abundant ions is advantageous when parent ions of the highest abundance are of primary interest (Bateman et al. 2014). In contrast, data-independent acquisition (DIA) protocols have less bias because the MS2 isolation windows are systematically stepped throughout a precursor m/z range such that all precursors in that m/z range are subjected to MS2 analysis, regardless of the precursor abundance (Egertson et al. 2015; Kawashima et al. 2019). While these data require more intensive deconvolution than DDA spectra, available software can deconvolute DIA spectra with high quantitative accuracy (Lou et al. 2023; Demichev et al. 2020).

We compared DDA and DIA for barcode identification by linking ∼25,000 v1 barcodes to muGFP for single-sample readouts that lack the peptide identification benefit of matching peptide IDs between runs. Whereas DIA used the MS2 signals for quantification, our DDA protocol relied upon the MS1 signal for this purpose. The unoptimized DIA protocol (at 500,000 resolution, 5µm packing silica) yielded more peptide identifications than our DDA protocol, supporting the hypothesis that DIA is more reliable in detection of these barcodes at higher pool complexities. We further improved sensitivity and quantitative accuracy using columns with finer particle sizes (from 5µm to 1.9µm) that provided higher chromatographic resolution and reducing the orbitrap MS1 resolution (from 500,000 to 30,000) to permit a more even survey of the sample. Together these resulted in a doubling of the barcode identification rate (30% to 60%, Figure 1d).

Despite the increase in peptide identification rate from our optimized DIA protocol, we still observed 40% barcode dropout. Barcodes with a high hydrophilicity score, and particularly barcodes rich in acidic residues Asp and Glu, were detected at lower rates (Supplementary Figure 1). Using this information, we generated a v2 barcode set with fewer acidic residues. LC elution times for 95% of detected barcodes were accurately predicted within 0.5 iRT (Figure 1c), confirming that Prosit predictions generalize with high accuracy to synthetic peptide sequences, and this library achieved an overall 86.6% barcode recovery rate with the optimized DIA protocol (Figure 1d). Further, the high detection rate observed amongst these barcodes through our purification protocol suggests that vast majority of these barcodes do not impact soluble expression of muGF and are, therefore, suitable for screening libraries of proteins for properties related to soluble expression (see barcode criterion 3 above).

We investigated whether barcode pre-assignment could reduce the high rates of barcode swapping (∼25%) observed by Egloff and colleagues during PCR amplification steps and explained as mega-primer formation due to high sequence homology. We sought to reduce barcode swapping by maximizing coding sequence distance over the set of proteins being examined, maintaining the same coding DNA sequence for each barcoded variant of the same protein. For our single oligo libraries (those not requiring two-oligo assembly), 99% of barcodes exclusively mapped to the design of interest, and for the two-oligo assembly, requiring an additional qPCR amplification to stitch together 5’ and 3’ oligos from separately amplified subpools using homologous junctions unique to each design, 75% of barcodes exclusively mapped to the intended design (Figure 1e). The low rate of barcode mismapping considerably increases library coverage, enabling screening of larger libraries.

### Screening beta barrel monomers

We applied our improved v1 mass spectrometry barcoding approach to a set of *de novo*-designed small beta barrels with six different barrel topologies, four of which were previously identified as protease-resistant via yeast surface display (Kim et al. 2023). Motivated by the observation that many of the designs were either insoluble or not monomeric when expressed individually (likely due to off-target strand pairing interactions), we set out to use size exclusion chromatography (SEC) to identify well-behaved designs from a set of individually pre-characterized controls (104 designs, not screened for protease resistance) and protease-resistant designs (416 designs) (Figure 2a).

**Figure 2:**
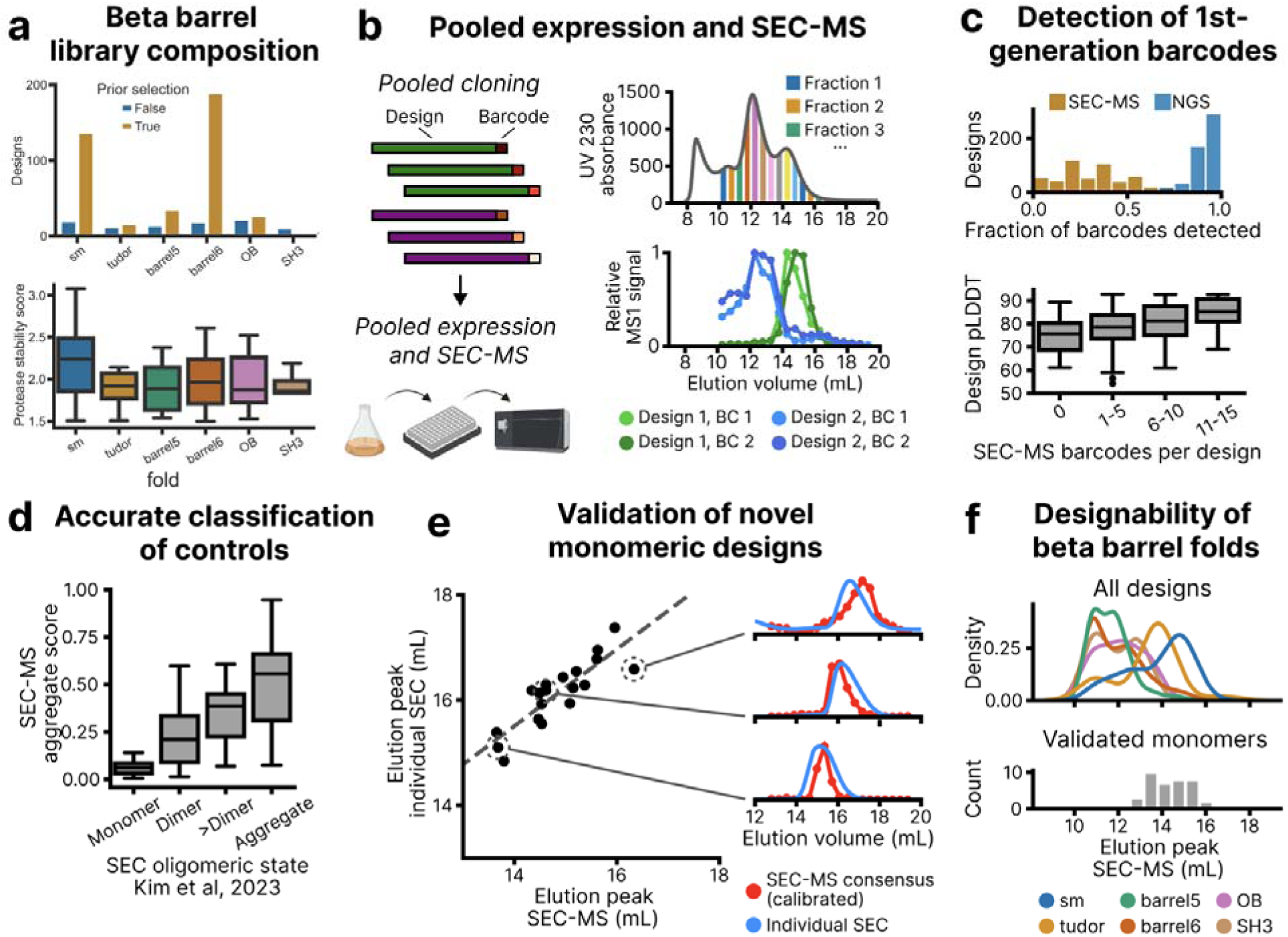
Evaluation of barcoding approach on library of designed monomer beta barrels. **(a)** Library distribution. Most designs were pre-selected for protease stability via yeast display (true and false in panel legend). An additional 104 controls were chosen solely by *in silico* metrics and individually characterized separately. **(b)** Designs and cognate barcodes were co-synthesized on oligonucleotide arrays. The library was cloned, expressed, and purified as a single pooled sample. After fractionation by SEC (top), barcodes were purified and quantified from each fraction by LC-MS/MS. Relative barcode abundance across fractions was used to reconstruct SEC traces (bottom). **(c)** Most barcodes were detected via next-generation sequencing (NGS), whereas a substantial fraction of barcodes were not detected in SEC-MS analysis (top). Barcode detection rate in SEC-MS was correlated with design quality measured by AlphaFold pLDDT (bottom). **(d)** For control designs individually characterized in a prior publication, aggregate score (fraction of consensus eluting between 10 and 12 mL) was predictive of off-target oligomeric state (all beta barrels were designed to be monomeric). **(e)** Novel monomeric designs were individually expressed without barcodes and characterized by SEC. Elution peaks agreed well between the pooled SEC-MS and individual SEC, up to a fixed offset (dashed line; added as a fixed global offset to SEC-MS consensus in examples on the right). **(f)** Comparison of designs grouped by fold (top) to validated monomers used as internal library controls (bottom) identified highly designable beta barrel folds.

An oligonucleotide library was synthesized with a median of 18 randomly pre-assigned v1 barcodes per design (total of 520 designs; 9,080 barcodes, ranging from 9-18 barcodes per design) (Figure 2a, Supplementary Figure 2a). Deep sequencing the *E. coli* library prior to protein expression confirmed high sequence fidelity; 98.9% of reads with exact barcode sequences also contained the exact intended design sequence (Figure 1e, Supplementary Figure 2b). The NGS results indicate a low frequency of barcode-design mismapping, which validates barcode pre-assignment as a strategy to streamline the library preparation process.

Pooled expression, Ni-NTA purification, SEC fractionation, and barcode purification resulted in 21 samples which were analyzed by LC-MS/MS (Figure 2b). A total of 2804 barcodes were detected accounting for 504 designs, with all 2804 barcodes detected in at least 5 samples (median 5 barcodes per design) (Figure 2c, Supplementary Figure 2c & 2d). The distribution of barcodes detected per design was highly skewed and correlated with design quality measured by AlphaFold pLDDT, with 30% of designs accounting for 50% of detected barcodes (Figure 2c, Supplementary Figure 2e), in part due to insoluble designs not being represented in SEC.

We reconstructed a per-barcode elution profile using the relative abundance of barcodes across SEC fractions. After removing barcode elution profiles with fewer than 5 data points, we defined a consensus elution profile for each design, normalized to the column volume (Figure 2d, Methods). In most cases, individual barcodes for the same design showed highly similar elution profiles, with elution peaks of individual barcodes having <10% coefficient of variation for 81% of designs, suggesting that most barcodes are non-perturbative to design behavior. (Supplementary Figure 2f). We took the fraction of the elution profile within the expected range for a protein aggregate as an aggregate score; this led to a well-defined separation between control designs previously reported to be monomer, dimer, or aggregate based on individual SEC characterization (Figure 2d; all designs were intended to be monomeric, so those that formed dimers are also more likely to have aggregate species). A subset of designs exhibited high variation among barcodes (Supplementary Figure 3). Given the high degree of correlation of barcode traces among validated well-behaved designs and the mechanism of barcode detection dropout, it is unlikely that the barcodes themselves are the source of design instability. Instead, we hypothesize that these may be marginally stable designs and hence more perturbed by barcode attachment than stably-folded designs.

To determine if pooled MS barcoding can identify successful designs, we selected 19 designs for individual characterization that showed consensus elution peaks in the expected range for monomers or dimers and low variation among barcode elution profiles (Figure 2e). For 16 designs we observed high soluble expression, with SEC elution peaks correlated strongly between pooled and individual expression (Figure 2d, Supplementary Figure 4). Notably, 16/19 (84%) screen hits designed using the SH3, barrel6, or barrel5, and OB beta barrel topologies eluted within 1 mL of the expected monomer elution volumes, whereas these designs showed a success rate of only 13/24 when selected solely based on protease resistance (Kim, et al. 2023). Consistent with these results, we found minimal correlation (r^2^ = 0.08) between protease stability score and either pooled SEC elution peak or number of barcodes detected (Figure 2e). Further structural characterization is described in Kim et al 2023.

We observed that success rates for monomeric design varied across six fold families, four native (1) oligosaccharide-binding – OB, 2) Src homology 3 – SH3, and two of SH3 subclasses: 3) small barrel – sm (Youkharibache et al. 2019), and 4) Tudor domains (Lasko 2010), and two *de novo* scaffolds, 5) b5 and 6) b6, previously described (Lasko 2010; Kim et al. 2023). Of these six families, barrels with sm folds had a higher monomer success rate than all other families (Figure 2f). On the other hand, non-Tudor/non-sm SH3, OB, b5, and b6 barrels tended to adopt higher order oligomers. Many of the Tudor barrels eluted as dimers, consistent with previous observations that Tudor domains can form homodimers (Tong et al. 2015; Cui et al. 2012; Zhao et al. 2007).

### Screening hallucinated oligomers

Encouraged by the ability to identify successful monomeric designs, we next applied pooled MS barcoding to a ∼10x larger library of 4,495 oligomers. The assembly of oligomers in solution is not well recapitulated in yeast display, so library-scale methods have not been applied to designed oligomers to date. For this design library, we expected successful designs to form highly stable complexes due to their large oligomeric interfaces. These complexes first assemble in the clonal environment of individual cells, and if stable should be maintained through cell lysis, pooled purification, and pooled SEC (Figure 3a). We selected oligomers with a protomer size comparable to the beta barrel library (65 amino acids) designed via a recently described hallucination method (Wicky et al. 2022). Of note, most (but not all) of the designs exhibit cyclic symmetry, so we opt for using C-notation herein to describe the number of subunits.

**Figure 3:**
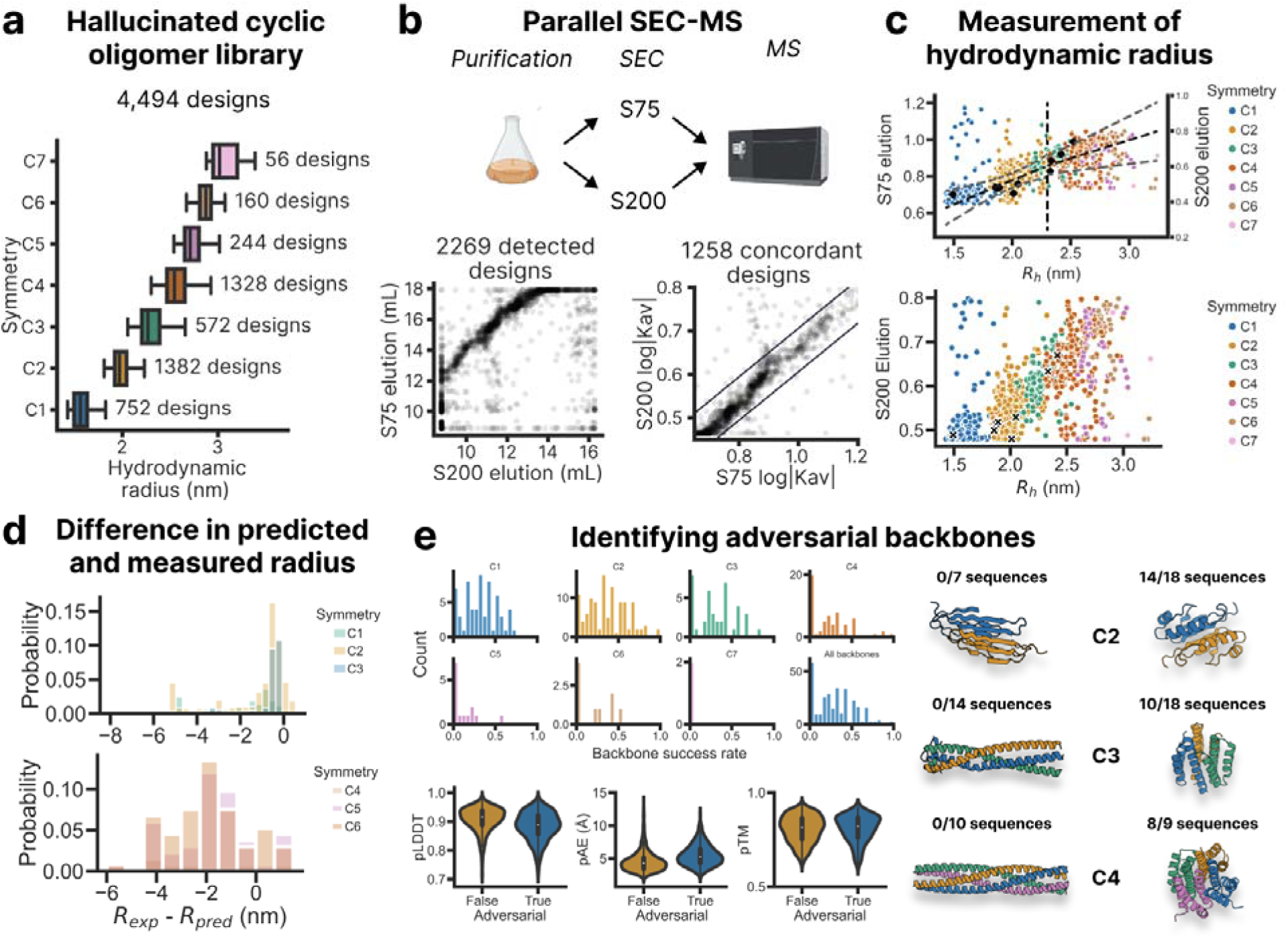
Characterization of small cyclic oligomers. **(a)** Cyclic homo-oligomers were designed by AlphaFold hallucination of backbone geometry with a symmetric loss function followed by sequence design with ProteinMPNN (top). Up to 18 sequences were designed per backbone in order to investigate the contribution of backbone quality to design success (bottom). **(b)** The library was fractionated using S75 and S200 SEC columns in order to capture a wide range of sizes. Sizing across the two runs was consistent for a majority of detected designs (bottom left). Designs were declared concordant if their normalized retention time (Kav) followed the expected linear relationship (bottom right). **(c)** For many concordant designs, rescaled Kav was proportional to hydrodynamic radius predicted from the design model. Shown in black are controls individually validated by SEC in published work. **(d)** Difference between the experimental and predicted radius of concordant designs for different symmetries. **(e)** A per-backbone failure rate was defined as the ratio of concordant designs to designs detected by NGS for each parent backbone (top). Adversarial backbones, defined as backbones with failure rate >0.85, had AlphaFold metrics indistinguishable from non-adversarial backbones (bottom) despite dramatically different success rates (examples on right).

A library of 4,495 designs was synthesized, including 82 controls from the beta barrel monomer library, 127 hallucinated controls published with detailed experimental characterization, and 4,286 new hallucinated designs (Figure 3a, Supplementary Figure 5a), ranging from C1 to C7 (22,475 barcodes, 5 barcodes per design). Sequencing quality control, pooled expression, purification, SEC, and MS barcode analysis were performed as for the beta barrel monomer library (Supplementary Figure 5), except that two separate SEC-MS runs were performed with S75 and S200 columns to capture the full range of elution volumes. After filtering for barcodes detected in at least 5 SEC samples, we obtained 5,200 barcodes representing 2,909 designs (23.14% barcodes representing 63.11% of ordered designs, Supplementary Figure 5). Of these 2,909 designs, 1,130 (39.81% of the library) had peak elution volume coefficients of variation <10% (Supplementary Figure 5e & 5j).

A calibration curve measuring elution volumes of proteins previously characterized by SEC, SEC-MALS, and/or crystallography was constructed to predict the elution volume of library members as a function of molecular weight and hydrodynamic radius (Figure 3b). The filtered dataset was then categorized based on whether the observed SEC-MS elution volume was within a bootstrapped 95% confidence interval, corresponding to the fraction with the highest MS1 area. Out of the 2,909 designs, 43% had elution volumes that were predicted by the calibration curve (Figure 3b, Supplementary Figure 6a). We defined success at the level of designs based on agreement between observed and predicted elution volumes, and at the level of individual backbones based on whether the backbone gave rise to at least one successful design. Compared to monomer design, which might fail due to (1) lack of soluble expression or (2) aggregation, the design of higher symmetries is further complicated by two additional failure modes: (3) lack of assembly or (4) formation of off-target assemblies. At the SEC stage, we observed aggregation failure modes for all cyclic oligomers: for C1 designs (designed monomers), an asymmetric bimodal distribution indicated that most C1 designs eluted at either the expected elution volume or in the void as an aggregate (Supplementary Figure 6a). For C2+ designs, we observed trimodal distributions, with densities observed a) where C1 designs elute, b) at the expected elution volume, and c) in the void where aggregates elute (Supplementary Figure 6). Higher order symmetries C4+ with MWs approaching 40 kDa had major peaks that were poorly resolved from the void peak on the S75 column, so we analyzed the elution profiles for these symmetries on an S200, which has greater resolution in the 30-100 kDa range (Figure 3c). Here, C4+ symmetries were better differentiated from the void volume, and we observed broad overlapping peaks for designed C4-6 oligomers. Given the oligomers have similar molecular weights and diameters, this result suggests a multiplicity of unintended oligomeric states for these designs. The difference in closing angle between C3, C4, and C5 is greater than the difference in closing angle between C6, C7, and C8, and hence obtaining specificity for the intended oligomerization state is more difficult for higher cyclic symmetries (Edman et al. 2023). Indeed, within each oligomer class, the success rate decreased as symmetry number increased, from 64.62% for C1s (n = 506) to 34.00% for C5 (n = 100) (Figure 3d, Figure 3e, Supplementary Figure 6c; C6 was an exception). The AF2 prediction metrics pAE and pTM (which were not used to curate the input design set) were not significantly different between successful and unsuccessful designs (Figure 3e), but many of the unsuccessful designs were somewhat less compact than backbones with high success rates (Figure 3e, right).

### Screening large helical oligomers

We sought to apply pooled MS barcoding to larger proteins by using oligo assembly to circumvent the length limits of pooled oligonucleotide synthesis. Cost-effective oligo pools currently have a maximum length of 300 nt, which constrains designs with pre-assigned barcodes to ≤74 aa to permit padding with subpools specific adapter sequences. Pairwise oligo assembly enables pre-assigned barcoding of much larger designs (≤154 aa), at the cost of decreased sequence representation due to differences in assembly efficiency among designs, and decreased sequence fidelity due to chimeric off-target assemblies. Chimeric assemblies are particularly concerning for peptide-based mass spectrometry, which relies on short sub-sequences as proxies for full-length proteins. To mitigate these issues, we developed a sequence design pipeline that minimized DNA homology among designs and used the same reverse translation for all barcoded variants of the same design, so assembly complexity scales with the number of designs but not the number of barcodes (Figure 4a).

**Figure 4:**
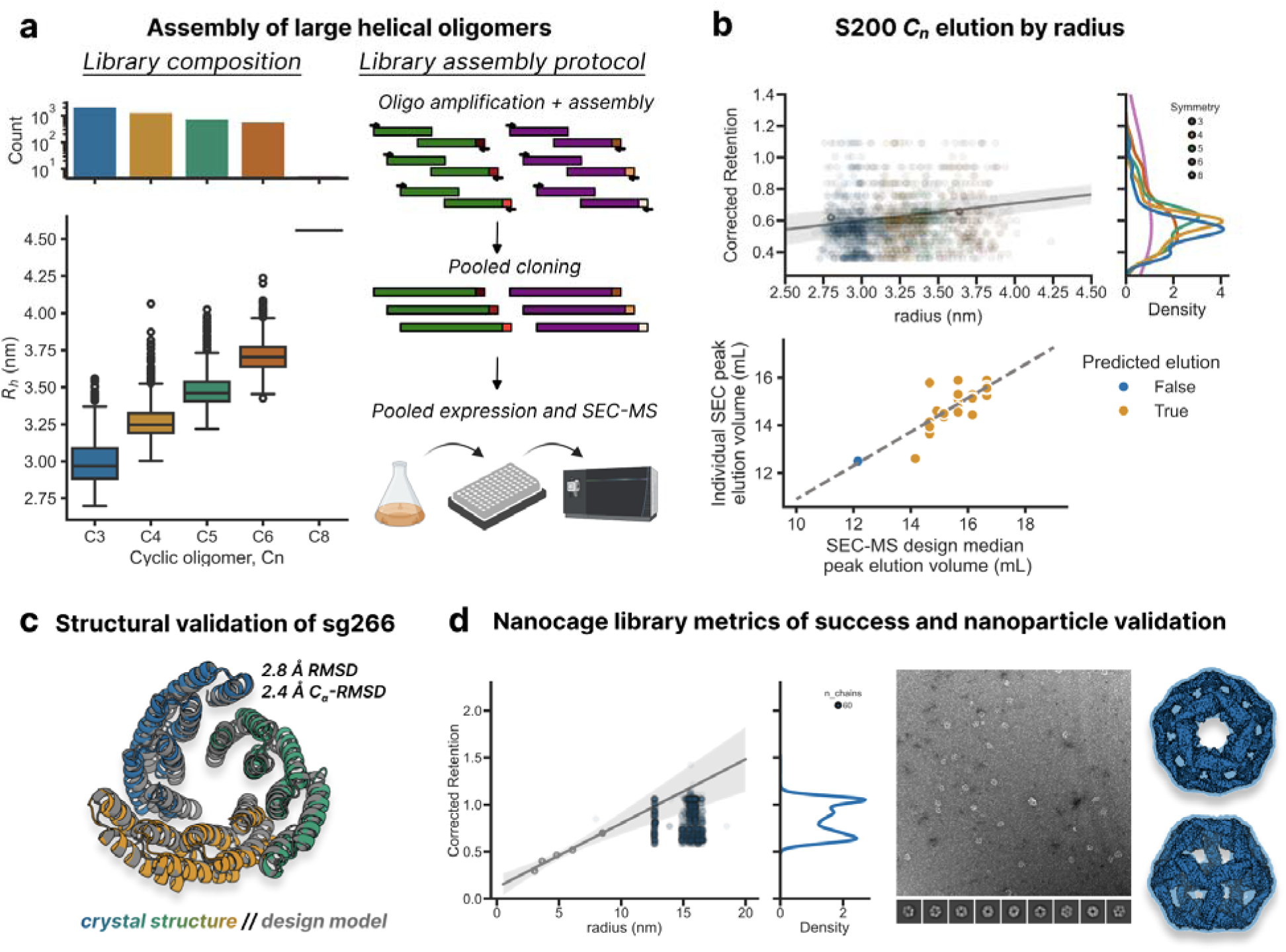
Characterization of libraries of large oligomer designs. **(a)** A library of designed large cyclic helical oligomers (>74 aa) of varying hydrodynamic radius with symmetries ranging from C3-C8 was assigned barcodes prior to chip-based DNA oligo synthesis (left). Owing to their large size, DNA oligos were subjected to assembly in pooled format, requiring a unique homologous sequence between part A (5’ fragment) and part B (3’ fragment) oligos to minimize formation of chimera sequences with mis-assigned barcodes. **(b)** Cyclic symmetries were fractionated on an S200, which permitted separation across the range of assembly states (top). Designs elected for follow-up were clonally expressed and purified at 50mL scale without barcodes, and their peak elution volumes on an S200 (bottom, y-axis) were compared to peak volumes from pooled expression (bottom, x-axis). **(c)** Crystal structure of sg266 (blue, green, and orange), a helical trimer hit from the screen, overlain with its design model. **(d)** Pooled elution profile of the I3 nanocage library (left). Negative stain electron micrograph (middle) and cryo-EM structure (right) of a hit from the screen, I3-08.

To test pooled MS barcoding of larger proteins, we barcoded a set of 5,068 homo-oligomers designed from curved helical repeats, ranging in symmetry from C2 to C6 (26,805 barcodes, minimum 5 barcodes per design, design length 119-156 aa). Deep sequencing after *E. coli* transformation showed 77.8% of barcodes covering 92.8% of all designs, with a median of 5 barcodes detected per design (Supplementary Figure 7). Of all barcoded designs, 99.4% of barcodes correctly and uniquely mapped to the design of interest, determining that the intended barcode pre-assignment was maintained throughout pooled DNA assembly. Pooled expression, purification, SEC, and MS barcode analysis were performed as for the beta barrel monomer library, except that SEC conditions were adjusted for an Superdex 200 10/300 GL column to optimize separation of oligomers. After filtering for barcodes detected in at least 5 SEC samples, we obtained 8311 barcodes (3060 designs, 60.38% of all designs), 52.1% of which had peak coefficients of variation < 10%.

We selected 28 designs with ≥2 barcodes detected, low peak coefficient of variation (<10%) among barcodes, and elution peaks within the expected range for their given symmetry and radius for individual characterization (Figure 4b). 26/28 designs showed high soluble expression, and SEC elution peaks correlated strongly between pooled and individual expression (Supplementary Figure 7). Overall, 27 (96.1%) of these designs eluted within the 95% confidence interval of the standard curve calibrated on hydrodynamic radius, and 15 of these designs (57.7%) eluted within 1 mL of the expected elution volume (Supplementary Figure 8). We selected one of these successful oligomers, sg266, to carry forward into crystallographic studies. We determined sg266 to adopt the modeled trimeric state by x-ray crystallography with an RMSD of 1.36Å between design model and the solved structure (Figure 4c). Additional structural characterization for these designs are published in (Gerben et al. 2023).

We further increased the size range to ∼215 kDa in a SEC-MS screen of one-component I3 nanoparticles interfaces with constant scaffolds (n=1173 cages, with 4 barcodes per design). The library was designed to interrogate interface metrics for assembly of icosahedral particles with identical trimer frameworks and fully redesigned interfaces. This library was subjected to pooled expression, purification, and SEC, and subsequent MS barcode isolation and analysis from SEC fractions were performed to analyze barcode traces. Filtering for barcodes detected in at least 5 SEC samples, we obtained 2434 barcodes mapping to 1090 designs with 643 (56.4%) having barcode peak elution volume coefficients of variation <10% (Figure 4d). For validation, 12 designs demonstrating assembly in the library format were individually expressed alongside aggregated and unassembled designs (Supplementary Figure 10). One of these hits, I3-08, was taken forward for structural characterization (Figure 4e), which revealed it to be an icosahedron of the on-target state. Upon expression of individual clones at the 50mL scale, cages appeared to adopt multiple oligomerization states on SEC, with peak elution volumes agreeing with peak values observed on SEC-MS, suggesting the desired assembly state is most abundant. We reasoned that these nanoparticles have a propensity to aggregate at the high concentrations expected at clonal 50mL scale, but not observed at the SEC-MS 50mL expression scale, which contains a heterogeneous pool of mixed expression. Nonetheless, the predominance of a peak at the expected elution volume corroborates the utility of this assay in large nanocage assemblies en masse.

Collectively, these data demonstrate that the oligomerization states of thousands of protein assemblies ranging in size from 10 kDa to 200+ kDa can be resolved in a single SEC run.

### Evaluation of v2 barcode library on tetrahedral nanoparticles

The v2 barcoding library was developed to improve detection of barcodes to permit high-confidence identifications of hits in complex pools. To evaluate the utility of the v2 barcode library, we set out to screen a library of putative *de novo* tetrahedra (n = 1,187 designs of 80 aa each, 7 barcodes per design, 8309 total barcoded designs) for homogenous nanoparticle assembly (Figure 5a). RFDiffusion has been successful in generating novel folds and topologies (Dauparas et al. 2022; Watson et al. 2023), and deep learning based sequence design has enabled the design of highly soluble proteins (Dauparas et al. 2022). While a diverse array of structures can be generated with these new design methods, robust nanoparticles are challenging to identify because these multi-component assemblies require reversible assembly of multiple weak interfaces to ensure particle homogeneity (Wargacki et al. 2021). As such, we aimed to use our higher fidelity barcoding set to identify novel, highly soluble tetrahedral nanoparticles.

**Figure 5:**
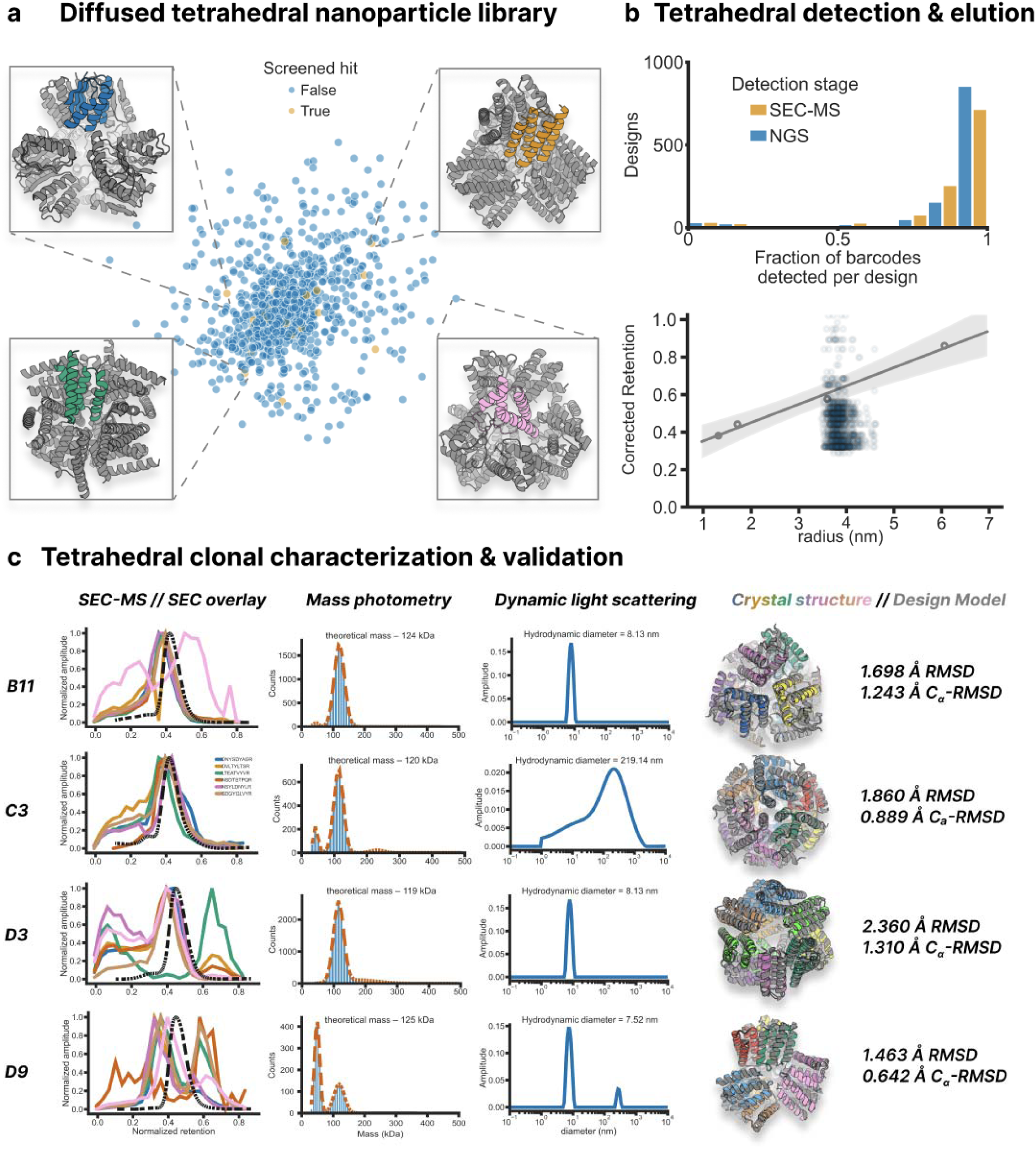
Tetrahedral nanoparticle characterization using 2nd generation barcodes. **(a)** A library of one-component tetrahedral backbones were generated using RFDiffusion and ProteinMPNN was used to design backbones. Sequences were then filtered by AlphaFold2 (pLDDT > 90) to predict the structure of the protomer (in color). Protomers were assessed for similarity by TM-score against all other library members. Multidimensional scaling of the all-by-all TM-score was plotted in 2D space. **(b)** Each library member was assigned 7 barcodes from the v2 barcoding set and ordered as a C-terminal genetic fusion as a single DNA oligo pool (n=8309 oligos). The DNA library was amplified, assembled, and transformed before being subjected to NGS and SEC-MS. Barcodes representing all 1,187 designs were detected at a comparable rate to NGS (top, median of 7, mean of 6.27 barcodes per design). The library was fractionated with an S200 SEC column in order to capture the expected range of assembly states. Designs were declared concordant if their corrected retention time (√(-log_10_(Kav)) followed the expected linear relationship (bottom). **(c)** Of the 13 hits identified by SEC-MS, four resulted in crystal structures. These four designs demonstrate concordance between the SEC-MS (color, solid), peak elution volumes and the clonally expressed (black, dashed) SEC trace at 50 mL scale (far left). Mass photometry (middle left) and dynamic light scattering (middle right) provide finer grained insight into the multiplicity of assembly states for C3 and D9 that is not evident by SEC. Crystal structures (far right) were solved to ≤3 Å resolution that corroborate the design model hypothesis.

From two ssDNA oligos per barcoded design, we used pairwise oligo assembly to construct full-length dsDNA encoding each design and a unique C-terminal barcode prior to cloning into a pET29b(+) plasmid encoding a C-terminal 6x-His tag. The library was then transformed, expressed, and purified prior to injection onto an S200 Increase 10/300 GL, as previously described for the large helical oligomers. NGS at 169x coverage identified 7429 correctly mapped barcodes (89.3% of all ordered barcodes) representing 1141 designs (96.1% of all ordered designs, Supplementary Figure 11). Of all detected barcodes on NGS, 80.9% exclusively mapped to the intended design (Figure 1e, Supplementary Figure 11). Barcodes were purified from 250µL SEC fractions and spectra were acquired with the optimized DIA LC-MS/MS acquisition protocol (Methods). Corroborating the results from the muGFP optimization study, 8003/8309 (97.3%) of barcodes were detected pre-filtering, and 98.3% of designs had ≥5 barcodes detected per design, with 48.4% of designs accounting for 50% of all detected barcodes (Supplementary Figure 12).

Peak elution volumes for all barcodes for a given design displayed low overall variation, with 81.3% of designs demonstrating CV < 10%, validating that the v2 barcode set is mostly well tolerated. Using a standard curve constructed from peak elution volumes of globular standards (Methods), 14 hits with peak elution volume CV < 10% that fell within the 95% CI of the standard curve for radius and/or MW (Methods, Supplementary Figure 13) were selected for follow-up expression and purification at 50mL scale with non-barcoded clones. Of these 14 designs, 13/14 expressed and 9/14 were monodisperse by SEC. Fractions at the expected nanoparticle elution volume were collected and subjected to complex sizing by mass photometry and hydrodynamic diameter measurement by dynamic light scattering (Figure 5c). Of the 9 hits, 8/9 had peaks corresponding to the expected molecular weight of a 12-mer assembly, and 8/9 had predominant peaks on DLS at the expected hydrodynamic diameter. In total, 3 of the 9 had monodisperse peaks by SEC, mass photometry (which exhibits bias towards incomplete assemblies), and DLS (which exhibits towards large assemblies), suggesting robust assemblies with no significant off-target or fractional assemblies (Figure 5c, Supplementary Figure 13, 14). Of the 9 hits, 4/9 diffracting structures were solved to ≤3 Å resolution (Figure 5c). The structures were very similar to the design models for each of these designs with ≤ 3 Å all-atom root-mean-square deviation to the design model across all 12 chains, corroborating the design hypotheses for these cages.

Overall, the v2 barcoding set increases the barcode detection rate from 30% to 90% which greatly enables identification of rare complex assemblies in a diverse structural pool.

## Discussion

We were able to substantially increase the power of the peptide barcoding coupled to mass spectrometry approach introduced by Egloff, Seeger, and coworkers. Egloff, et al. used random peptide barcodes stochastically linked to a nanobody library derived from llama immunization to measure solution-phase properties. We extend this work in two key ways. First, we leveraged pooled oligo synthesis and assembly to generate libraries directly from *in silico* sequences up to 154 aa in length. We streamlined library construction by a) omitting cloning of the binder library into *E. coli* for pre-enrichment and b) attaching barcodes at the oligo level to circumvent barcode mapping via next-generation sequencing. Second, we explicitly designed barcode peptides to optimize mass spectrometry based detection and to minimize perturbation to attached proteins. Our approach should enable scaling the number of barcodes to up to 120,000 to greatly increase the number of proteins that can be analyzed simultaneously. Moving forward, recent advances in deep learning-based sequence design now enable elimination of possible interference from appended barcodes by embedding the barcode directly in the coding sequence.

Our mass spectrometry-barcoding approach enables large-scale testing of designed protein properties inaccessible to previous screening methods based on surface display and proxy reporters. Using this approach, we assessed the solubility and oligomerization state of thousands of computational designed proteins in parallel. Our results across diverse folds — including beta barrels, hallucinated cyclic oligomers, large helical cyclic oligomers, and protein nanoparticles — suggest that, despite potential liabilities of pooling (such as cross-design interaction and low concentration of each design), designs with the desired properties (e.g., intended oligomerization state) can be robustly identified. Data generated by large-scale testing can provide feedback on the design process, exemplified by our analysis of systematic failure modes in cyclic oligomers generated by hallucination and deep learning-based sequence design. While we focused here on solubility and size-separation, pooled MS barcoding generalizes to virtually any protein property amenable to solution-phase fractionation, such as binding, folding stability, and permeability across biological barriers.

## Materials and Methods

*N.B. – all buffers prepared in Milli-Q-grade distilled deionized water (ddH_2_O) unless otherwise specified*

### Single oligo library cloning

Subpools were amplified from an oligonucleotide pool (Twist) with subpool-specific primers using qPCR with KAPA HiFi HotStart ReadyMix (Roche) and EvaGreen Dye (Biotium). To reduce template switching that could scramble the design-barcode linkage, a minimal PCR cycle count was determined for each sample via a test reaction. Amplified oligos were size-selected with a 2% agarose E-gel EX (ThermoFisher), quantified using a Qubit dsDNA Quantification Assay (Thermo), and cloned into a modified pET28b plasmid using NEBridge Golden Gate Assembly Kit (BsaI-HF v2) (New England Biolabs). Approximately 100 ng of assembled plasmid was transformed into *E. cloni* Express BL21(DE3) Electrocompetent Cells (Lucigen). After recovery for 1h at 37°C, cells were transferred to a 50 mL LB overnight culture with kanamycin selection, from which glycerol stocks were prepared. Transformation efficiency was assessed by counting CFUs from a 1:50,000 dilution of the recovery culture on LB-Kan100 plates (Teknova).

Primer sequences:

>DF_Nterm_pT05_fwd_1

CTACTGGTCTCaCAGTCGA

>DF_Cterm_pT05_rvr_1

ACTGGAGACGGTCTCaGTTA

>DF_Cterm_pT05_fwd_2

GACACGGTCTCtCAGTCGA

>DF_Cterm_pT05_rvr2

ACGATTCTGGGTCTCtGTTA

>DF_Cterm_pT09_fwd_1

GCTCTCGGTCTCgTACCATG

>DF_Cterm_pT09_rvr_1

AGATGGACTGGTCTCgTGCG

>DF_Cterm_pT09_fwd_2

CTATCATTCGGTCTCcTACC

>DF_Cterm_pT09_rvr_2

AGTCTAATTGGTCTCcTGCG

>oDF-287_foldit_fwd_0_NGS_tP5

TCCCTACACGACGCTCTTCCGATCTctactggtctcacagtcga

>oDF-288_foldit_rev_0_NGS_tP7

GTTCAGACGTGTGCTCTTCCGATCTactggagacggtctcagtta

>oDF-289_foldit_fwd_1_NGS_tP5

TCCCTACACGACGCTCTTCCGATCTgacacggtctctcagtcga

>oDF-290_foldit_rev_1_NGS_tP7

GTTCAGACGTGTGCTCTTCCGATCTacgattctgggtctctgtta

>DF_pT15_1_short_fwd

GGGTCTAGggtctcaAGGA

>DF_pT15_1_short_rvr

AGTACTCGggtctcaCGCT

#### Adapter sequences

Adapter sequences are terminal sequences that flank the DNA of the protein of interest. They are used to selectively amplify subpools from pooled single stranded oligo DNA. Their order on a sequence is:

5’ end adapter – coding sequence of protein of interest – 3’ end adapter.

##### Adapters for beta barrels (pair with DF_pT09 primers)

>DF_Cterm_pT09_1

5’ end – GCTCTCGGTCTCgTACCATG

3’ end – CGCAcGAGACCAGTCCATCT

##### Adapters for small cyclic oligomers (pair with DF_pT15_1_short primers

>DF_pT15_1

5’ end – TGCTTTGGGTCTAGggtctcaAGGA

3’ end – AGCGtgagaccCGAGTACTTCTGGT

>DF_pT15_2

5’end – GACATGATCTAGAGggtctcaAGGA 3’ end – AGCGtgagaccTCCGACCATTCTTT

##### Adapters for large helical oligomers (pair with)

##### Adapters for nanocages (pair with)

>jason_01_A

5’ end – GCGACTAGGGGTATGCTG

3’ end – AAgcttacgggcaacATGG

>jason_01_B

5’end –GCGAcccactggcataaTT

3’ end – gcctgtgcgaaacATGG

>jason_02_A

5’ end – GCGAcgtgaccaccaagg

3’ end – AAcccagtaagggtcATGG

>jason_02_B

5’ end – GCGAgcgaccttagagtTT

3’ end – cagagaggtcagcATGG

>jason_03_A

5’ end – GCGAagtcccttaccctt

3’ end –AAgctcccgtatcagATGG

>jason_03_B

5’ end – GCGAgagggtctacagaTT

3’ end – ctcggtccacgatATGG

### Two-oligo assembly library cloning

The previously described protocol for two-oligo assembly (Klein et al. 2016) mirrored the single oligo library construction protocol with two exceptions. First, chip oligo ssDNA for 5’ and 3’ fragments was amplified with KAPA HiFi HotStart Uracil+ReadyMix (Roche) along with uracil-containing adapters on the inner termini (3’ end of the 5’ fragment and 5’ end of the 3’ fragment). After gel extraction, fragments were resuspended in 20µL ddH_2_O and incubated with 2 µL USER Enzyme (New England BioLabs) in a thermocycler at 37°C for 15 minutes followed by 22°C for 15 minutes. After cooling, the entire USER digest mixture was mixed with 10 µL 10x NEBNext End Repair Enzyme Mix (New England BioLabs), 5 µL NEBNext End Repair Reaction Buffer (New England BioLabs), and 63 µL ddH_2_O for incubation on a thermocycler at 20°C for 30 minutes. DNA was then cleaned up and prepared for qPCR amplification with the primers to the outer adapters (5’ end of the 5’ fragment and 3’ end of the 3’ fragment) and KAPA HiFi Hotstart *without uracil*. The rest of the protocol is reflected in the single oligo library construction protocol, where the amplicon running at the combined length of the 5’ + 3’ fragments is extracted, cleaned up, and cloned into pET28b(+).

### Next Generation Sequencing

Plasmid DNA was extracted from overnight cultures using a ZymoPURE II Plasmid Midiprep Kit (Zymogen). The cloned insert containing design and barcode was PCR-amplified and size-selected using the same protocol as for library cloning. Sample barcodes and Illumina sequencing adapters were added in a second qPCR, and paired-end 2×300 nt reads were acquired on MiSeq (lllumina).

### Protein purification and size-exclusion chromatography (SEC)

Glycerol stocks (100µL) were used to inoculate 50mL Studier’s Autoinduction media (Studier 2014) in 250 mL baffled flasks. Cultures were grown for 20h at 37 °C, and cell pellets were harvested via centrifugation (10 min, 4000 x *g*). Pellets were resuspended in 25 mL lysis buffer (25 mM Tris HCl pH 8, 300 mM NaCl, 40 mM Imidazole, 1 mM DNase I, 10 µg /mL lysozyme) and lysed by sonication (Q500 Sonicator Dual Horn ¾ “ probes, Qsonica, 5 min, 85% amplitude, 15s on/off cycles). Lysate was clarified by centrifugation (14,000 x *g*, 20 min.). The pellet was resuspended in 1 mL of SEC running buffer (50 mM Tris HCl pH 8, 150 mM NaCl) to create the “insoluble fraction” sample, while the supernatant was defined as the “soluble fraction” sample. Ni-NTA resin (Qiagen) was used to isolate His-tagged proteins from the soluble fraction. Specifically, wash buffer (25 mM Tris HCl pH 8, 300 mM NaCl, 40 mM imidazole) and then elution buffer (25 mM Tris HCl pH 8, 150 mM NaCl, 400 mM imidazole) were applied to the resin. Libraries were eluted in 2 mL, of which 100 µL was defined as the “injection” sample, and 1 mL was used for SEC. SEC was performed on an equilibrated S75 Increase 10/300 GL or S200 Increase 10/300 GL at a flow rate of 0.8 mL/min in SEC running buffer. Fractions were collected in 0.25 mL intervals from 8 mL to 20 mL (Wicky et al. 2022).

### Crystal Structure Determination

Crystals were produced using the sitting drop vapor diffusion method. Drops with volumes of 200 nL in ratios of 1:1, 2:1, and 1:2 (protein:crystallization) were set up in 96-well plates at 20 °C, using the Mosquito from SPT Labtech. Drops were monitored using the JANSi UVEX imaging system.

For sg266, diffraction-quality crystals appeared in a mixture of 0.2 M Ammonium sulfate, 0.1 M HEPES pH 7.5, 25% w/v Polyethylene glycol 3,350.

Diffraction quality crystals appeared in 0.1 M Sodium chloride, 1.6 M ammonium sulfate and 0.1 M Sodium HEPES pH 7.5 for B11; 10 % v/v PEG 400, 0.05 M MES pH 6, 0.1 M Potassium chloride and 2 mM Magnesium chloride hexahydrate for C3; 1.4 M Sodium malonate dibasic monohydrate pH 7.0 for D3 and 21 % w/v PEG 3350, 0.1 M MES pH 6.0 and 0.15 M Sodium chloride for D9.

Crystals were cryoprotected before being flash frozen in liquid nitrogen before being shipped for data collection at synchrotron. Data collection was performed with synchrotron radiation at the Advanced Light Source (ALS) on beamline 8.2.2/8.2.1 or NSLS2 on beam line FMX/AMX.

X-ray intensities and data reduction were evaluated and integrated using either XDS (Kabsch 2010) and merged and scaled using Pointless and Aimless in the CCP4 program suite (Winn et al. 2011). Structure determination and refinement starting phases were obtained by molecular replacement using Phaser (McCoy et al. 2007) using the design model for the structures.

Following molecular replacement, the models were improved using Phenix autobuild (McCoy et al. 2007; Adams et al. 2010); efforts were made to reduce model bias by setting rebuild-in-place to false and using simulated annealing. Structures were refined in Phenix (Adams et al. 2010). Model building was performed using COOT (Emsley and Cowtan 2004). The final model was evaluated using MolProbity (Emsley and Cowtan 2004; Williams et al. 2018). Data collection and refinement statistics are available in Supplementary Table 1. Data deposition, atomic coordinates, and structure factors reported in this paper have been deposited in the PDB 8VEA, 9DE9, 9DEA, 9DEB, 9DEC.

### Barcode design

*Barcodes consisting of 8-12 variable amino acids were appended to the N or C-terminus of designed amino acid sequences prior to reverse translation and pooled, single stranded oligo DNA synthesis. Each sequence was tagged with multiple barcodes to average out barcode-specific perturbations. Barcodes were designed as KxxxxR, to enable tryptic digest would yield a doubly-charged precursor of the form xxxxR. Notably the xxxx regions omitted certain amino acids: F & W to minimize hydrophobicity, C & M to minimize barcode cross-reactivity, H, K, & R to avoid alternate charge states greater than +2 during positive mode ionization. In the v2 barcoding set, the number of D & E was restricted to be ≤ 2 to minimize the number of alternate charge states. Barcode sequences were selected to be orthogonal by high resolution LC-MS/MS, with a minimum chromatographic separation of 8 iRT units (predicted using Prosit (Gessulat et al. 2019), and a minimum MS1 mass-to-charge separation of 10 parts per million. Sequences containing designs and appended barcodes were reverse translated as in the yeast surface protease assay, except for specifying E. coli codon usage, and ordered as a 300 nt oligo pool (Twist Bioscience)*.

### Barcode Isolation

For each sample, 100µL of sample was added to 100 µL of Lys-C buffer (8M urea, 100mM Tris HCl, pH 8) plus 1 µg of Endoproteinase LysC (New England Biolabs). Samples were incubated in a shaker at 37°C for 4-6 hours. To isolate His-tagged barcodes, 15 µL of Dynabeads™ His-Tag Isolation and Pulldown (Thermo) was added to each sample, incubated at 25°C for 5 minutes, and a magnetic rack was used to separate beads from supernatant. Beads for each sample were washed twice with 200 µL of barcode wash buffer (50 mM Tris HCl, pH 8, 150 mM NaCl, 0.1% Tween-20), followed by two 300µL washes with trypsin buffer (50 mM Tris HCl, pH 8). Beads were then resuspended in 50µL trypsin buffer plus 0.25 μg trypsin Trypsin-ultra (New England Biolabs) and incubated at 37°C overnight for at least 6 hours shaking at >1000 rpm.

### Mass Spectrometry

NanoLC analytical columns were prepared with capillary with 75um inner diameter, cut to 40 cm with laser puller, and packed with ReproSil-Pur 120 C18Aq media 5µm or 1.9µm silica (Dr. Maisch) at 1000 psi to a final column length of 15-20 cm. Trap columns were prepared with capillary with 75um inner diameter and cured overnight for polymer frit formation. Trap columns were packed at 500 psi with ReproSil-Pur 120 C18Aq media 5µm silica (Dr. Maisch) to a final column length of 1-3 cm. Columns were equilibrated on an EASY-nLC 1200 (Thermo). LC-MS/MS was run using an EASY-nLC 1200 and either an Orbitrap Fusion Lumos Tribrid (Thermo) or an Orbitrap Exploris 480 (Thermo) Mass Spectrometer at the University of Washington Proteomics Resource. Samples were prepared 50% in 0.1% trifluoroacetic acid, and 8µL of 1:2 diluted sample was loaded onto the trap columns at 2.5 µL/min before separation at 300 nL/min on the analytical column with an 89 min gradient (Solvent A = ddH_2_O with 0.1% formic acid, Solvent B = 80% acetonitrile with 0.1% formic acid, Gradient = 6-40% Solvent B). Peptides were ionized with nanospray ionization with a positive ion voltage of 2100 V at 300°C. Under the DDA protocol, MS data were acquired over 120 min with a cycle time of 3s utilizing the Thermo “Advanced peak determination” setting. MS1 scans were collected in profile with an Orbitrap resolution of 500,000 (480000 on the Exploris) and a precursor scan range of 450-900 m/z with the following parameters: RF lens – 30%, AGC target – Custom, Normalized AGC target – 175%, Polarity – positive. MS1 data went through MIPS (monoisotopic precursor selection) filtering for peptide peak determination, a charge state filter for 2-5 charge states, a dynamic exclusion filter (excludes precursor after n=1 times for 10 seconds, with a 10ppm low and high tolerance, and excludes isotopes) prior to MS2. Centroid MS2 data were acquired with isolation windows of 1.6 m/z and a normalized HCD collision energy of 27% at 15,000 resolution. MS1 scans collected under our optimized DIA protocol reduced MS1 resolution to 30,000.

Under the DIA protocol, samples are loaded at 2.5µL/min and run on a gradient at 400nL/min (Solvent A = ddH_2_O with 0.1% formic acid, Solvent B = 80% acetonitrile with 0.1% formic acid, Gradient = 6-40% Solvent B) over 89m. Nanospray ionization with positive spray voltage of 2000V, internal mass calibration of Easy-IC, expected peak width of 30s and charge state of 2 for MS1 settings. MS1 resolution w/ resolution of 30000 in a scan range 450-1000, RF lens% = 40, data collected in centroid. Standard AGC target, “automatic” maximum injection time mode, 1 microscan per scan. MS2 has an isolation window of 6m/z without isolation offset with a resolution of 15000 acquired over 150-2000m/z. Collision energy uses normalized HCD collision energy of 27%. RF lens% = 50. AGC target set for 1000% normalized AGC target with “auto” maximum injection time. 1 microscan is performed, and data is collected in centroid. The DIA window starts from 493.4742 and collects 61 spectra with a 6 m/z window to an end mass of 850.6366 and is under N loop control.

### In-silico design of tetrahedral nanoparticles

Tetrahedral nanoparticle backbones of 80AA per subunit were generated with RoseTTAFold Diffusion as previously described (Watson, et al.). Following diffusion, backbones were minimally downsampled, only filtering on external C-termini (for experimental purification) and inter-subunit contacts to ensure sufficient interface size. Backbones were then designed as homooligomers with ProteinMPNN at a sampling temperature of 0.1 and with 8 sequences per RFdiffusion-generated backbone (Dauparas, et al.). Candidate sequences were then predicted (asymmetric unit only) with AlphaFold2 (Jumper, et al.) and designs were filtered on pLDDT ≥ 85 and RMSD ≤ 1 Å, with 1,187 designs passing.

### Mass photometry

Mass photometry measurements were collected as previously described (Pillai et al. 2023) with a TwoMP (Refeyn) mass photometer. Pooled fractions corresponding to cage assembly were diluted 1:10,000 in SEC running buffer (50mM Tris, 300 mM NaCl, pH 8), and 10 µL dilute solution was applied to a gasket well for focusing and collection (1 minute) under a large field of view. Ratiometric values were converted to masses in kDa using known sizes of a 20 nM β-amylase standard in the same buffer (tetramer 224 kDa tetramer, 112 kDa dimer, and 56 kDa monomer). Expected masses were computed based on expressed protein sequence multiplied by 12, the number of subunits in a 1-component tetrahedron.

### Dynamic light scattering

Measurements were performed with standard settings for polydispersity and sizing with an UNcle (Unchained Labs), as previously described (Yang et al. 2024). To a glass cuvette, 8.8 µL of sample ∼1 mg/mL was applied, and data were collected at 25 °C with a 1s incubation time.

## Acknowledgments

Many thanks to the community of colleagues, friends, and scientists who’ve assisted with this work. Specifically, we thank Robert Ragotte, Matthias Glögl, and Samuel J. Pellock for thoughtful discussions and critical feedback. We thank Kandise VanWormer, Rafael Ticzon, Hernan Nuñez-Ortega, Ratika Krishnamurty, Kristina Herrera, Lance Stewart, and the myriad support personnel at the Institute for Protein Design who have provided an enabling environment to carry out this work. Various sources of funding supported this research for authors D.F., J.N.S., X.L., R.J., S.G., D.K., C.R., B.K., H.E., D.R.H., E.C.Y., B.I.M.W., L.F.M., A.K.B., A.K., E.B., E.J., B.S., J.M.L., I.G., D.V., A.A., L.S., M.J.M, & D.B: NSF Grant CHE-1629214 (A.K.B.), EMBO long-term fellowship ALTF 139-2018 (B.I.M.W.), Defense Threat Reduction Agency Grant HDTRA1-19-1-0003 (X.L., S.G.), National Institutes of Health’s National Institute on Aging, grant R01AG063845 (I.G), National Institutes of Health’s National Cancer Institute, grant R01CA240339 (I.G.), National Institutes of Health’s National Institute of Allergy and Infectious Disease, grant R0AI160052 (S.G., A.K.B.), DARPA program Harnessing Enzymatic Activity for Lifesaving Remedies (HEALR) under award HR0011-21-2-0012 (X.L. A.K.B., I.G., L.S.), National Institutes of Health’s National Institute on Aging, cooperative agreement U19AG065156 (D.R.H.), National Institutes of Health’s National Institute on Aging, grant U19AG065156 (D.R.H.), Spark Therapeutics / Computational Design of a Half Size Functional ABCA4 (B.I.M.W., L.S.), Defense Threat Reduction Agency Grant HDTRA1-21-1-0038 (I.G., D.V.), Juvenile Diabetes Research Foundation International (JDRF) grant # 2-SRA-2018-605-Q-R (X.L.), AMGEN & NovoNordisk (I.G.), Helmsley Charitable Trust Type 1 Diabetes (T1D) Program Grant # 2019PG-T1D026 (X.L.), Bill and Melinda Gates Foundation #OPP1156262 (X.L., S.G., A.K.), The Audacious Project at the Institute for Protein Design (J.N.S., A.K.B., D.R.H., L.S. S.G., D.E.K., C.R., B.K., E.C.Y., A.K., E.B., L.F.M.), The Nordstrom Barrier Institute for Protein Design Directors Fund (E.C.Y., I.G.), The Open Philanthropy Project Improving Protein Design Fund (S.G., C.R., B.I.M.W., A.K.B., E.J., E.B., I.G., L.S., D.B.), The Open Philanthropy Project Universal Flu Vaccine Fund (L.S.), Washington Research Foundation and Translational Research Fund (J.M.L.), Dr. Eric and Ms. Wendy Schmidt, and Schmidt Futures funding from Eric and Wendy Schmidt by recommendation of the Schmidt Futures program (J.M.L., I.G.), European Molecular Biology Organization (EMBO) Postdoctoral Fellowship (L.F.M.), & Howard Hughes Medical Institute (J.N.S., A.K.B., I.G., D.R.H. D.B.). Additionally, we want to thank the Advanced Light Source (ALS) beamline 8.2.2/8.2.2 at Lawrence Berkeley National Laboratory and National Synchrotron Light Source II for X-ray crystallography data collection. The Berkeley Center for Structural Biology is supported in part by the National Institutes of Health (NIH), National Institute of General Medical Sciences, and the Howard Hughes Medical Institute. The ALS is supported by the Director, Office of Science, Office of Basic Energy Sciences and US Department of Energy (DOE) (DE-AC02-05CH11231). At NSLSII the Center for Bio-Molecular Structure (CBMS) is primarily supported by the NIH-NIGMS through a Center Core P30 Grant (P30GM133893), and by the DOE Office of Biological and Environmental Research (KP1607011). NSLS2 is a U.S.DOE Office of Science User Facility operated under Contract No. DE-SC0012704. This publication resulted from the data collected using the beamtime obtained through NECAT BAG proposal # 311950 and # 313951.

