## Supplementary Information for "Massively parallel assessment of designed protein solution properties using mass spectrometry and peptide barcoding"

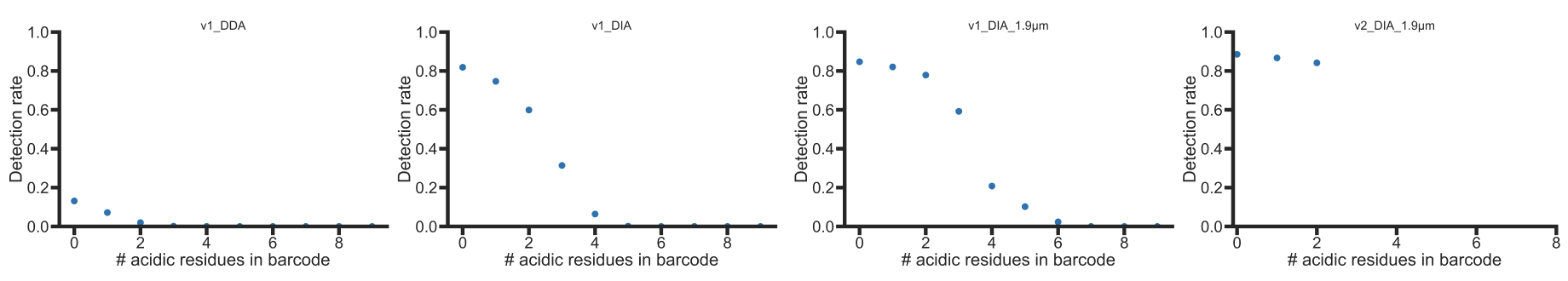

**Supplementary Figure 1.** Barcode detection rate by acidic residues across protocols using muGFP tagged with v1 (C-terminal) or v2 (N-terminal) barcodes. Barcode detection rate was quantified using the Skyline analysis pipeline for DDA or the DIA-NN analysis pipeline for DIA. The v2 library was restricted to < 3 acidic residues (Asp & Glu) to ensure higher detection rates.

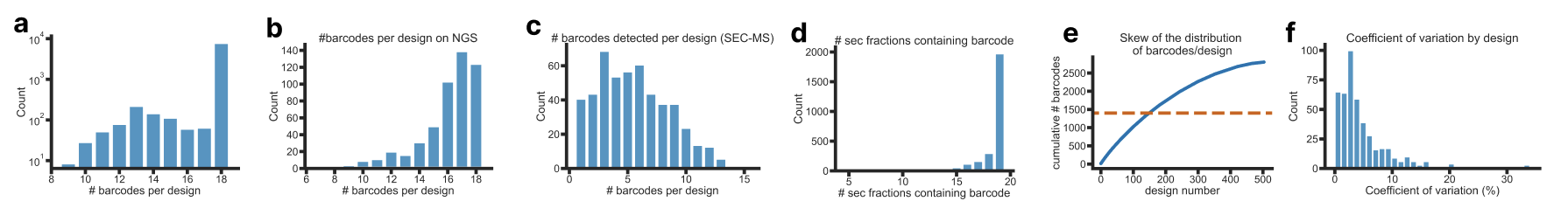

**Supplementary Figure 2.** Beta barrel library design and detection statistics. **(a)** distribution of barcodes ordered per design. **(b)** distribution of barcodes per design as detected by NGS. **(c)** distribution of barcodes per design as detected by SEC-MS. **(d)** distribution of barcode detection rate by SEC fraction. **(e)** skew of design representation based on SEC-MS barcode detection. 50% of the detected barcodes account for 30% of all designs. **(f)** coefficient of variation (CV) of barcode elution volume for SEC-MS. 81% of designs have CV < 10%.

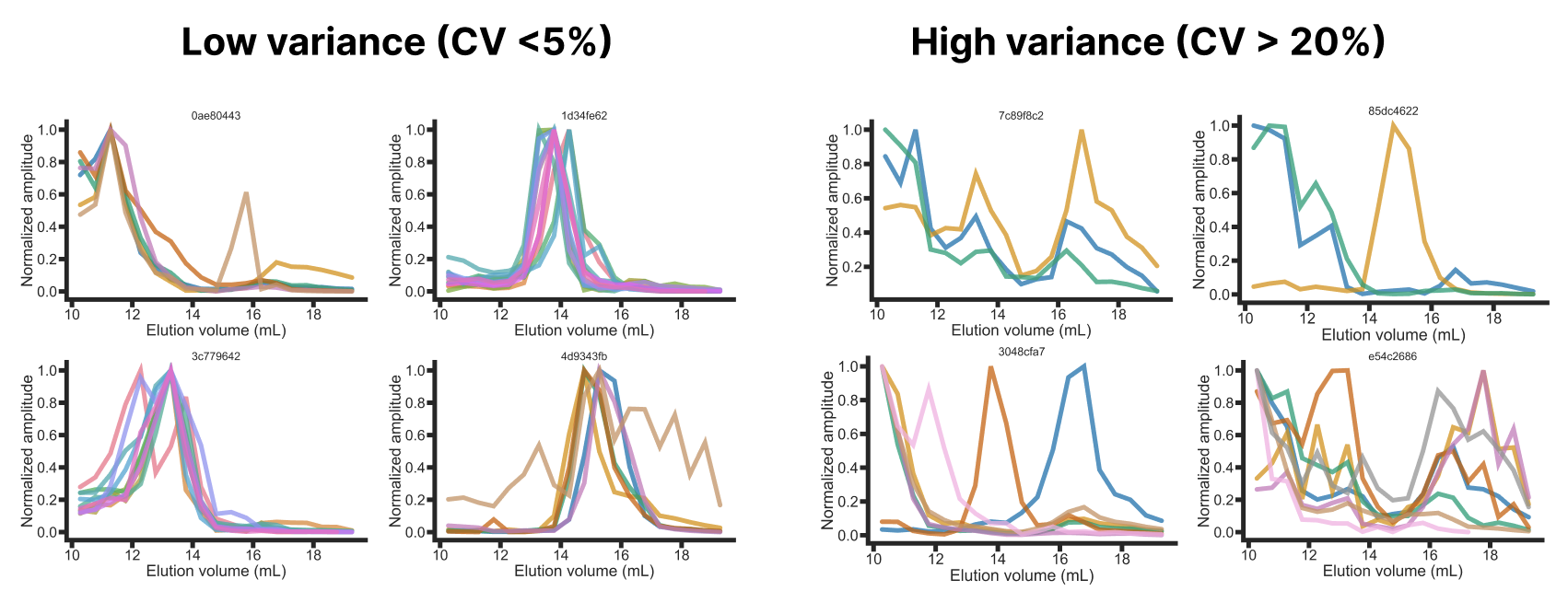

**Supplementary Figure 3.** Beta barrel library SEC-MS traces with low variability (< 5% coefficient of variation, CV) in peak elution volume (left) and high variability (>20%, right).

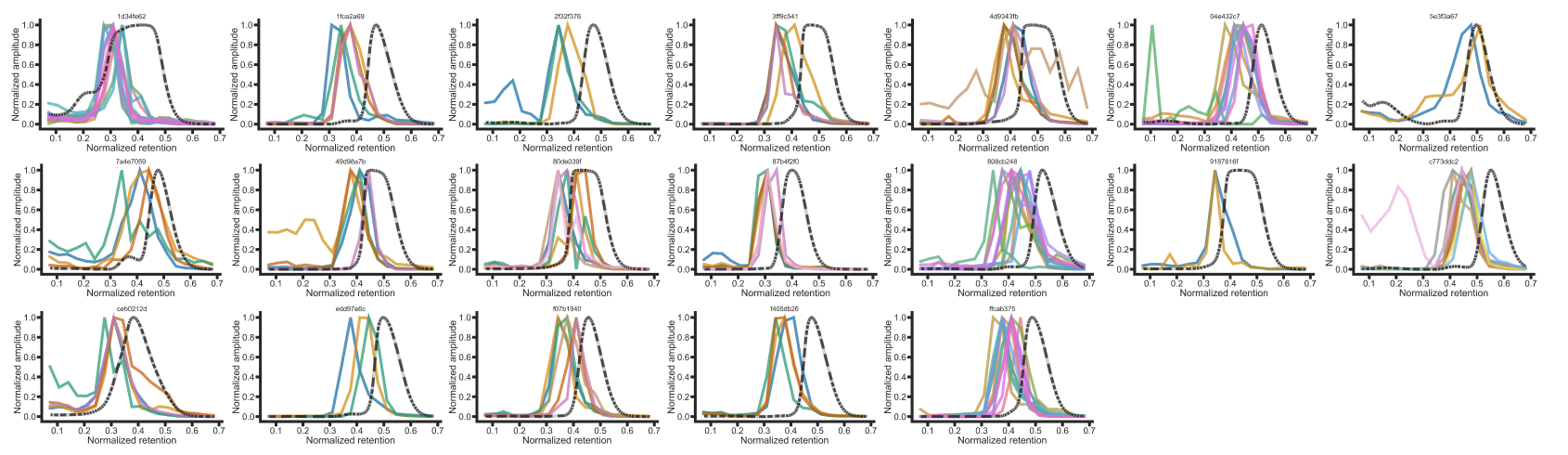

**Supplementary Figure 4.** Individual beta barrel SEC traces of validated monomers without barcodes (absorbance at 230nm, black dotted) overlaid with their corresponding barcoded SEC-MS traces (solid color).

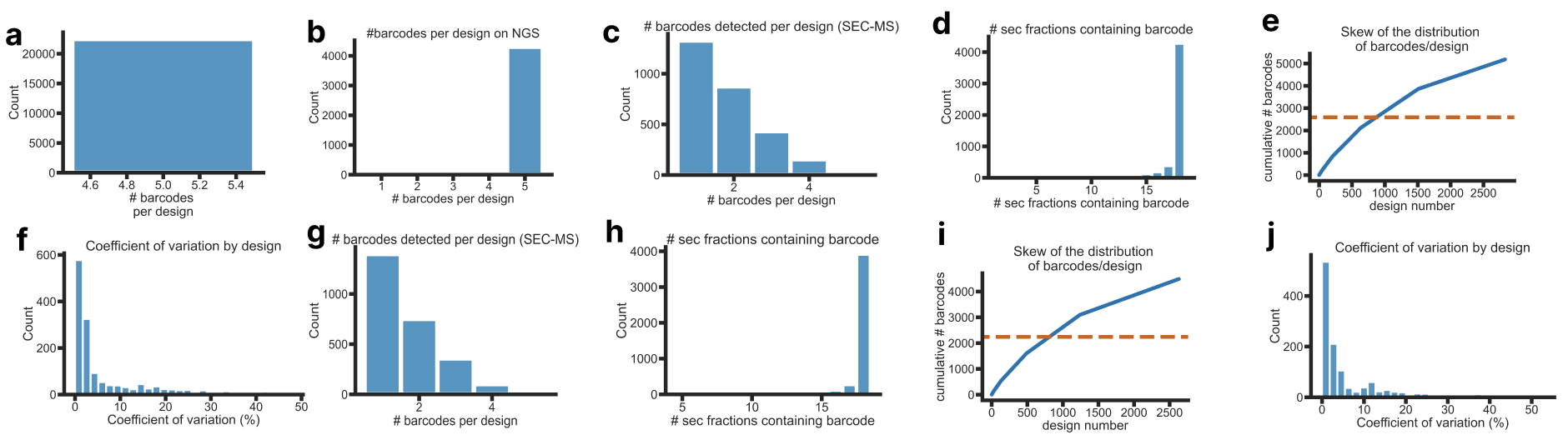

**Supplementary Figure 5.** Hallucinated cyclic oligomer design and detection statistics for S75 and S200 runs. **(a)** distribution of barcodes ordered per design. **(b)** distribution of barcodes per design as detected by NGS. **(c)** distribution of barcodes per design as detected by S75 SEC-MS. **(d)** distribution of barcode detection rate by S75 SEC fraction. **(e)** skew of design representation based on S75 SEC-MS barcode detection. 50% of the detected barcodes (orange, dotted) account for 29% of all designs. **(f)** coefficient of variation of barcode elution volume for S75 SEC-MS. **(g)** distribution of barcodes per design as detected by S200 SEC-MS. **(h)** distribution of barcode detection rate by S200 SEC fraction. **(i)** skew of design representation based on S200 SEC-MS barcode detection. 50% of the detected barcodes (orange, dotted) account for 29% of all designs. **(j)** coefficient of variation of barcode elution volume for S75 SEC-MS.

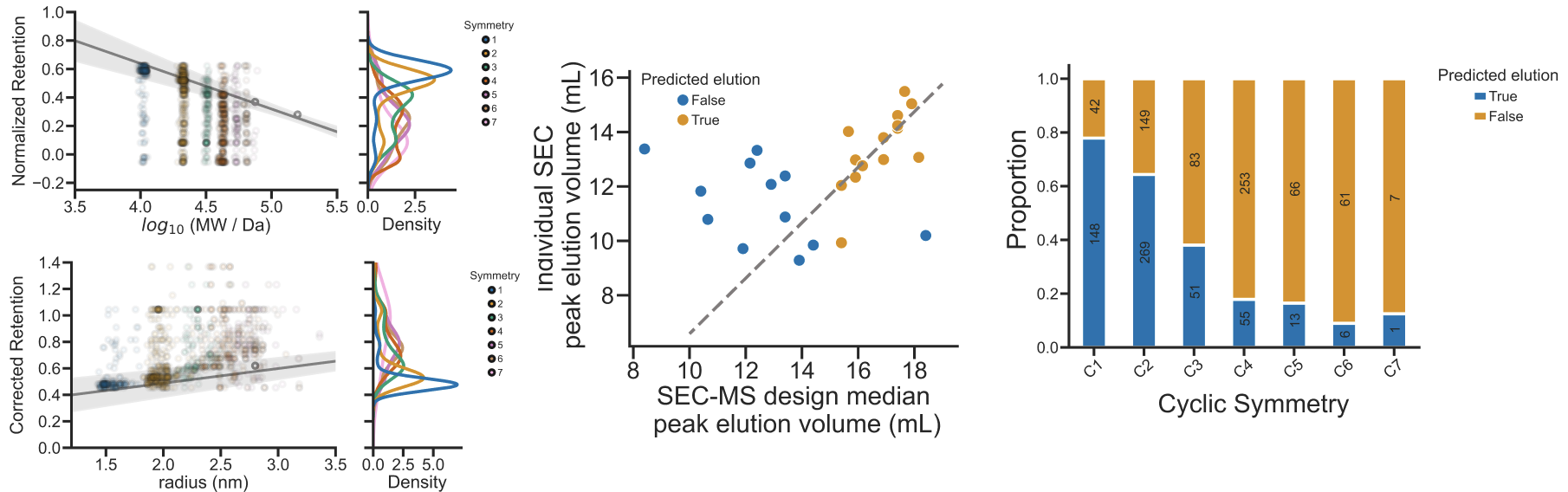

**Supplementary Figure 6. (a)** Hallucinated cyclic oligomer library SEC elution profiles for S75 (top) and S200 (bottom). **(b)** SEC peak elution volumes of hits expressed clonally at 50-mL culture scale (y-axis) vs barcode peak elution volume (x-axis) on an S200 column. **(c)** Design success rate based on concordance of predicted and measured elution volumes on SEC.

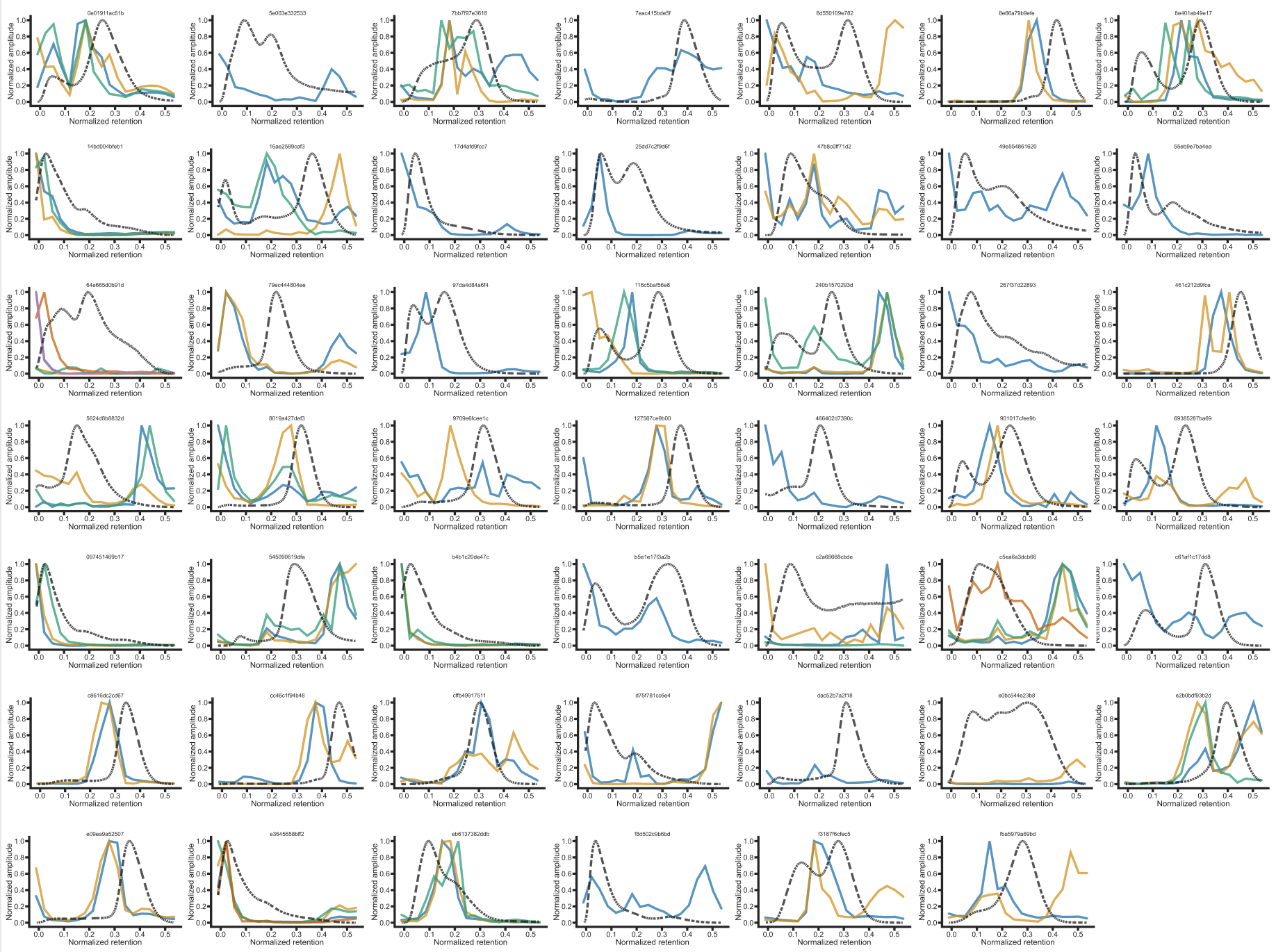

**Supplementary Figure 7.** Hallucinated oligomer hits S75 SEC-MS (solid color) overlaid with clonal SEC S75 data (Absorbance @ 280 nm, black dotted). Sequence-verified clones were expressed with a C-terminal 6x-His tag (without barcodes), purified via Ni-NTA, and sized on an S75 Increase 10/300 GL in 50 mM Tris, 150 mM NaCl, pH 8.

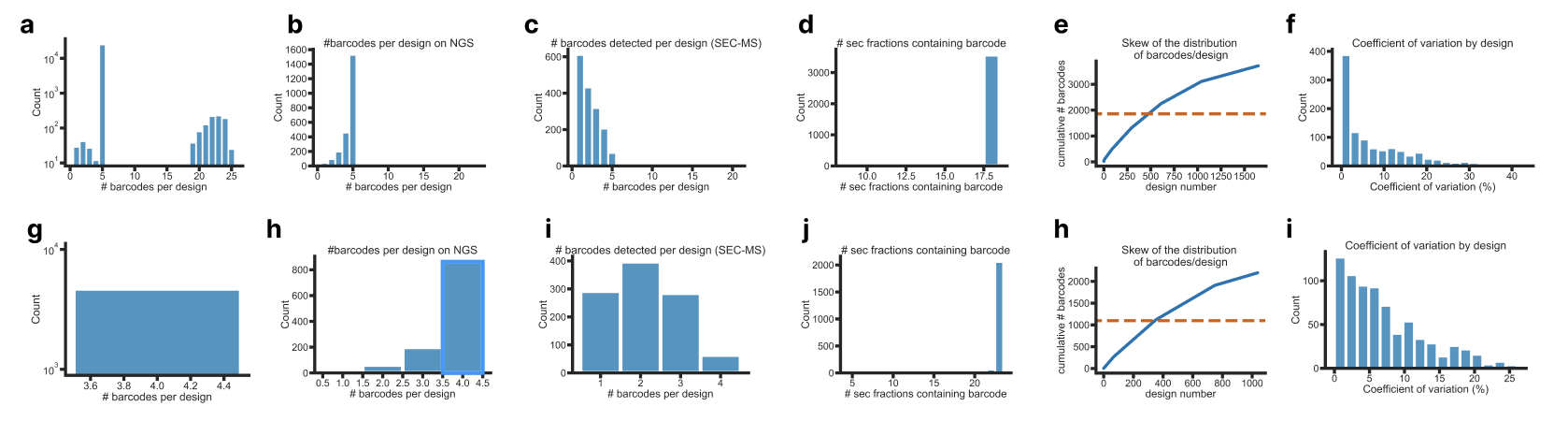

**Supplementary Figure 8.** Large helical oligomer **(top)** and I3 icosahedral nanocage **(bottom)** design and detection statistics. **(a & g)** distribution of barcodes ordered per design. **(b & h)** distribution of barcodes per design as detected by NGS. **(c and i)** distribution of barcodes per design as detected by SEC-MS **(d & j)** distribution of barcode detection rate by SEC fraction. **(e & k)** skew of design representation based on SEC-MS barcode detection. 50% of the detected barcodes account for 29% (top) and 33% (bottom) of all designs. **(f & i)** coefficient of variation (CV) of barcode elution volume for large helical oligomer (top) and I3 icosahedra (bottom) SEC-MS. 81% of large helical oligomer designs have CV < 10%, and 53% of I3 icosahedral designs have CV < 10%.

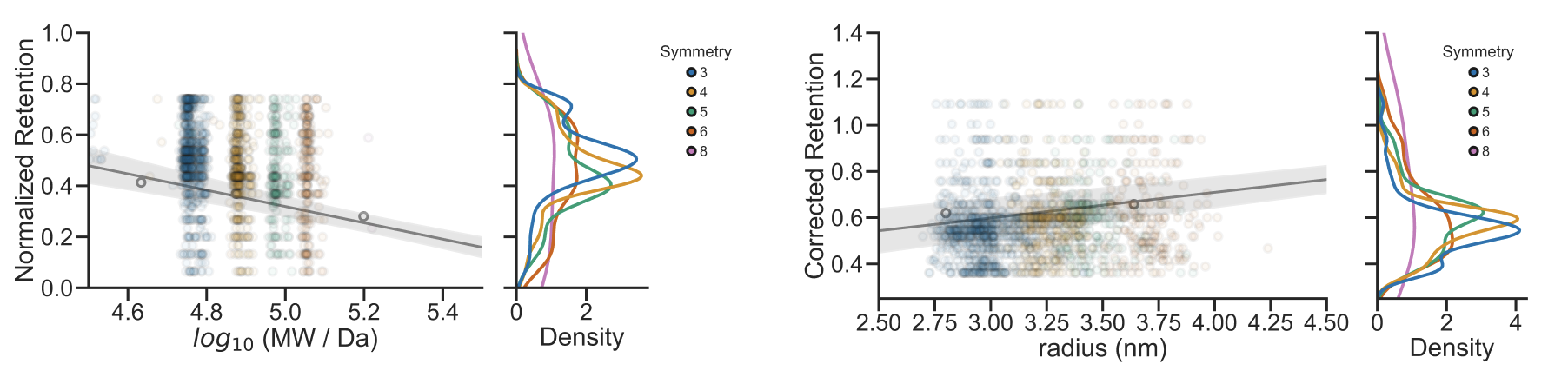

**Supplementary Figure 9.** S200 elution profiles of large helical oligomers along a standard curve. For each design, the median barcode elution profile was plotted vs either log_10_(molecular weight) in Da, or via computed hydrodynamic radius (see Supplementary Figure 11). Normalized retention (also known as K_av_) is based on the void volume of the S200 column (by elution of blue dextran, where K_av_ = 0) and the known column volume (24 mL – K_av_ = 1).

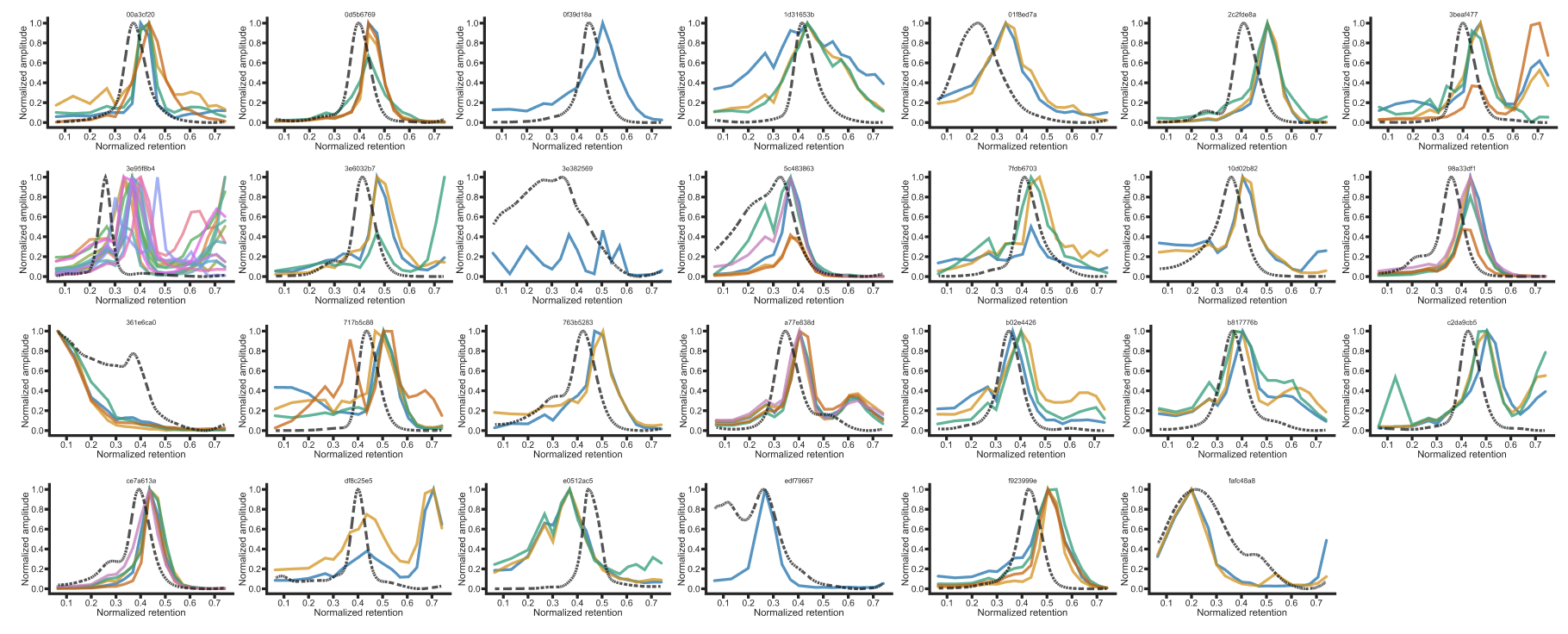

**Supplementary Figure 10.** Large helical oligomer hits SEC-MS overlaid (solid color) with clonal SEC data (Absorbance @ 280 nm, black dotted). Sequence-verified clones were expressed with a C-terminal 6x-His tag (without barcodes), purified via Ni-NTA, and sized on an S200 Increase 10/300 GL in 50 mM Tris, 300 mM NaCl, pH 8.

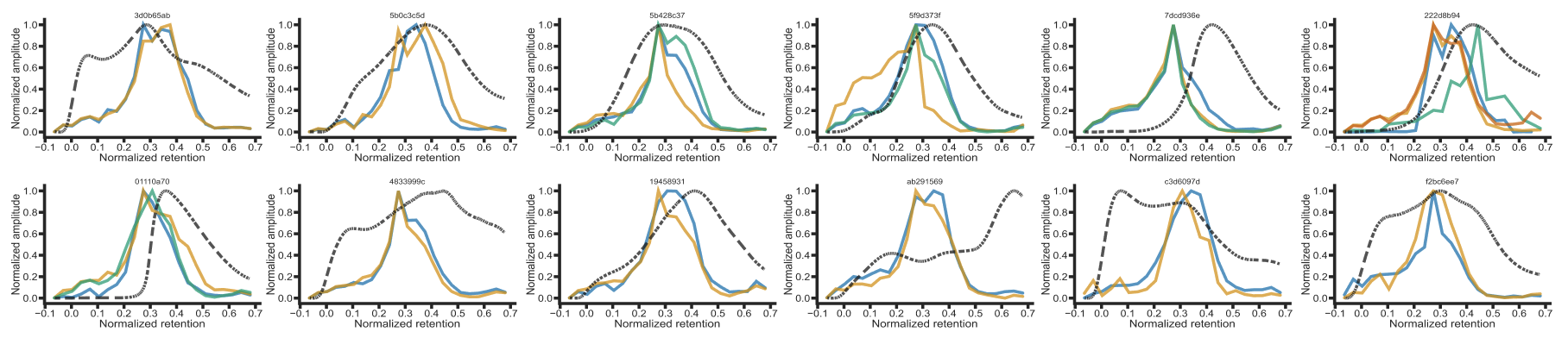

**Supplementary Figure 11.** I3 icosahedral hits SEC-MS overlaid (solid color) with clonal SEC data (Absorbance @ 280 nm, black dotted). Sequence-verified clones were expressed with a C-terminal 6x-His tag (without barcodes), purified via Ni-NTA, and sized on an S200 Increase 10/300 GL in 50 mM Tris, 300 mM NaCl, pH 8.

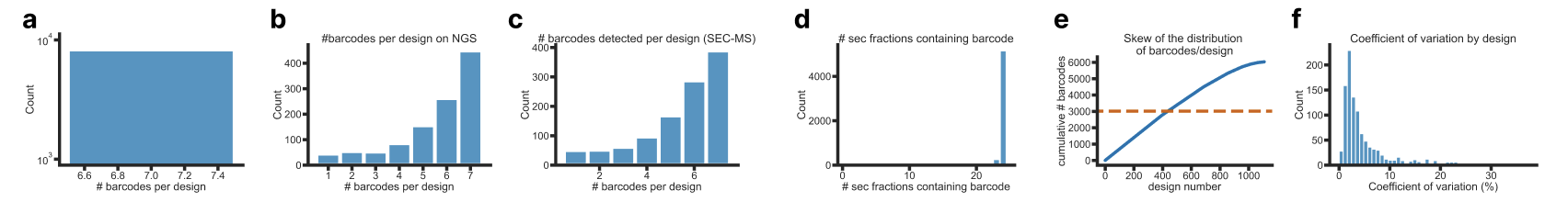

**Supplementary Figure 12.** Tetrahedra library design and detection statistics. **(a)** distribution of barcodes ordered per design. **(b)** distribution of barcodes per design as detected by NGS. **(c)** distribution of barcodes per design as detected by SEC-MS. **(d)** distribution of barcode detection rate by SEC fraction. **(e)** skew of design representation based on SEC-MS barcode detection. 50% of the detected barcodes account for 39% of all designs. **(f)** coefficient of variation (CV) of barcode elution volume for SEC-MS. 81% of designs have CV < 10%.

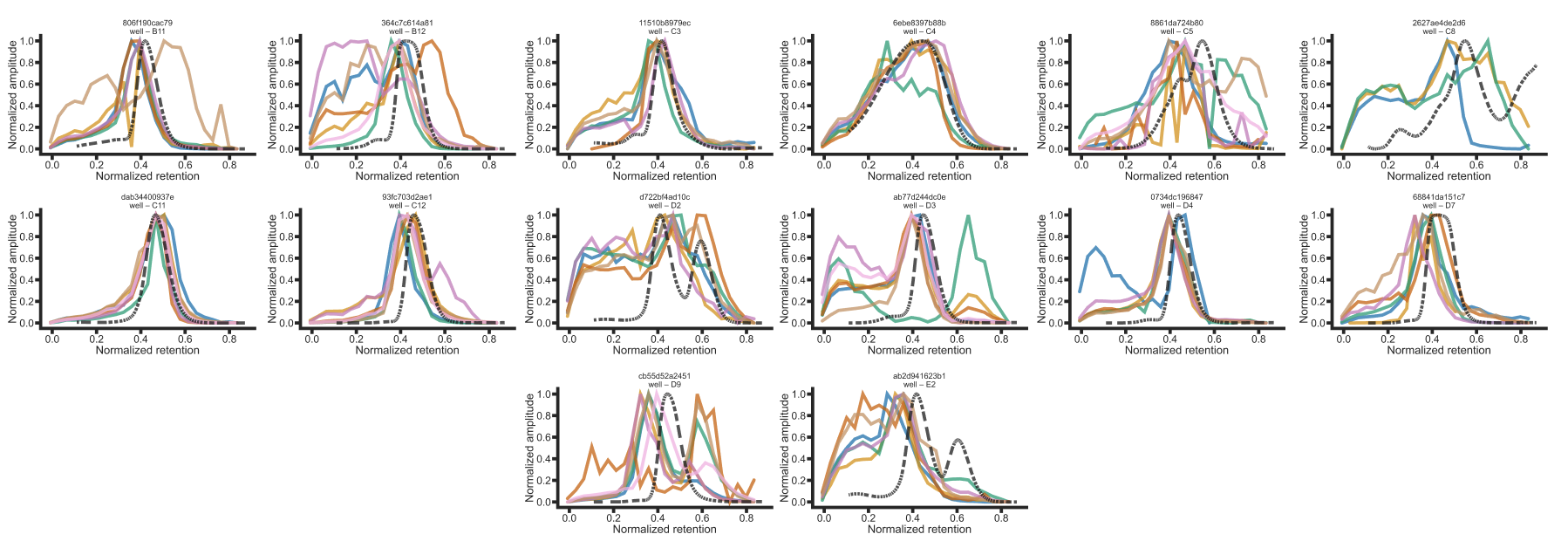

**Supplementary Figure 13.** Tetrahedral hits SEC-MS overlaid (solid color) with clonal SEC data (black dotted). Sequence-verified clones were expressed with a C-terminal 6x-His tag (without barcodes), purified via Ni-NTA, and sized on an S200 Increase 10/300 GL in 50 mM Tris, 300 mM NaCl, pH 8. Absorbance @ 230 nm was recorded as many of these proteins lack aromatics that absorb at 280 nm.

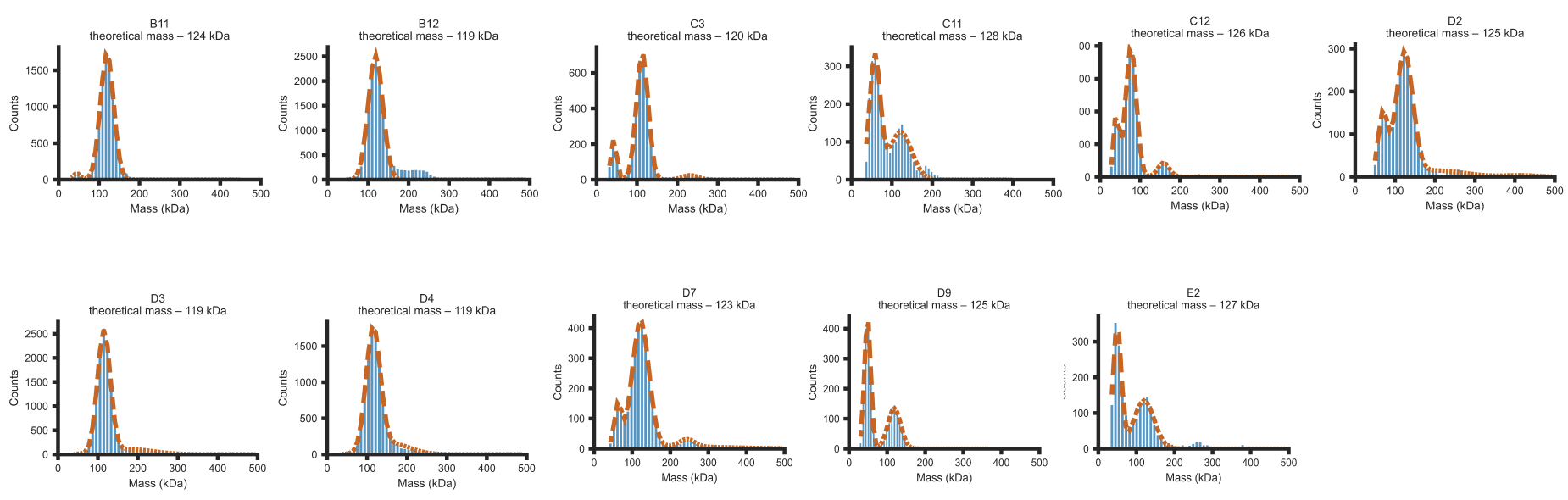

**Supplementary Figure 14.** Mass photometry data of hits from tetrahedral library. Sequence-verified clones were expressed, purified, and sized on an S200 in 50mM Tris, 300mM NaCl, pH 8. Peaks corresponding to expected elution volume were collected and 10 µL of 1:1000 diluted solution (in aforementioned SEC running buffer) was subjected to mass photometry analysis.

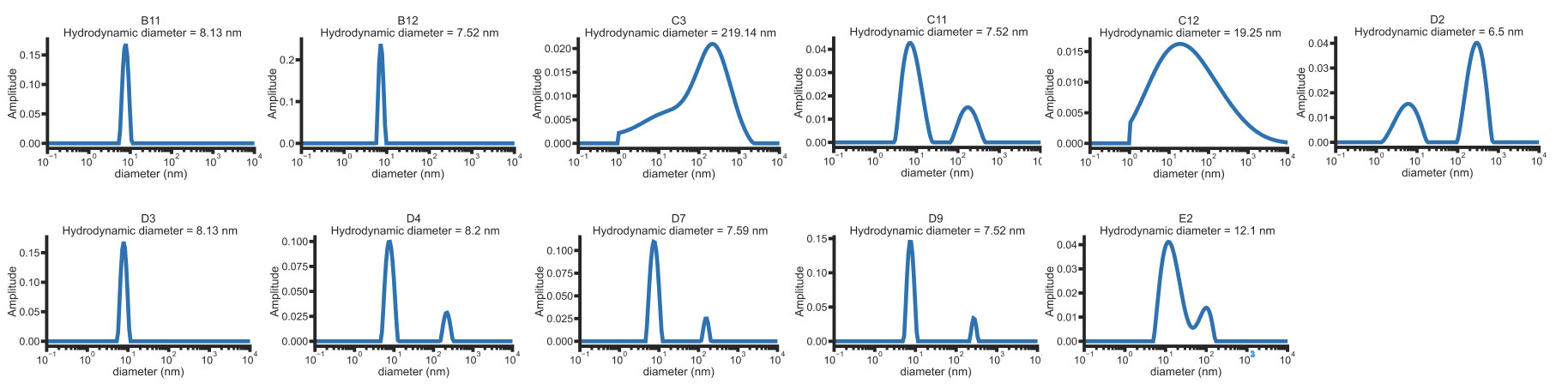

**Supplementary Figure 15.** Dynamic light scattering data of hits from tetrahedral library. Sequence-verified clones were expressed, purified, and sized on an S200 in 50mM Tris, 300mM NaCl, pH 8. Peaks corresponding to expected elution volume were collected and 8.8 µL of 1mg/mL solution to dynamic light scattering analysis.

**Supplementary Table 1.** X-ray diffraction data collection and refinement statistics

|  | SG266 (8VEA) | B11  (9DE9) | C3  (9DEA) | D3  (9DEB) | D9  (9DEC) |
| --- | --- | --- | --- | --- | --- |
| Resolution range | 41.58 - 3.30 (3.55 - 3.30) | 45.71 - 2.52 (2.62 - 2.52) | 36.32 - 2.87 (3.10 - 2.87) | 46.54 - 2.54 (2.60 - 2.54) | 31.86 - 2.04  (2.15 - 2.04) |
| Space group | *P 63* | *P 21 21 21* | *I 2 3* | *P 21* | *I 2 2 2* |
| Unit cell | 115.33, 115.33, 59.99; 90, 90, 120 | 45.71, 49.55, 76.02; 90, 90, 9 | 72.64, 72.64, 72.64; 90, 90, 90 | 75.03, 119.44, 116.12; 90, 91.62, 90 | 67.97, 69.27, 84.46; 90, 90, 90 |
| Unique reflections | 6977 (1381) | 6161 (654) | 1542 (305) | 306782 (5350) | 13040 (1879) |
| Multiplicity | 15.9 (14.7) | 12.8 (13) | 38 (38) | 3.4 (2.0) | 10.9 (11.2) |
| Completeness (%) | 99.69 (99.78) | 99.3 (97.8) | 100.00 (100.00) | 99.19 (99.05) | 100 (100) |
| Mean I/sigma (I) | 28.45 (10.97) | 14.8 (2.4) | 19.8 (2.8) | 9.0 (1.0) | 8.3 (1.2) |
| Wilson B-factor | 101.26 | 46.36 | 87.47 | 49.72 | 38.36 |
| R-merge | 0.074 (0.292) | 0.123 (1.065) | 0.172 (1.826) | 0.167 (2.056) | 0.159 (2.387) |
| R-pim | 0.018 (0.078) | 0.037 (0.315) | 0.029 (0.302) | 0.074 (0.900) | 0.053 (0.782) |
| CC_1/2_ | 1.00 (0.987) | 0.999 (0.866) | 0.999 (0.810) | 0.998 (0.363) | 0.996 (0.416) |
| Reflections used in refinement | 6977 (1381) | 5752 (1148) | 1539 (1539) | 67006 (4788) | 11792 (1220) |
| R-work | 0.1850 (0.2742) | 0.2650 (0.3202) | 0.2376 (0.2376) | 0.2131 (0.3106) | 0.2220 (0.3138) |
| R-free | 0.2423 (0.3231) | 0.3051 (0.3978) | 0.2722 (0.2722) | 0.2624 (0.3693) | 0.2781 (0.3671) |
| Number of non-hydrogen atoms | 3492 | 1283 | 609 | 14856 | 1921 |
| macromolecules | 3492 | 1261 | 609 | 14848 | 1906 |
| Solent | n/a | 22 | n/a | 8 | 15 |
| Protein residues | 429 | 159 | 79 | 1979 | 239 |
| RMS (bonds) | 0.004 | 0.003 | 0.004 | 0.002 | 0.002 |
| RMS (angles) | 0.64 | 0.549 | 0.70 | 0.49 | 0.46 |
| Ramachandran favored (%) | 96.93 | 96.77 | 93.51 | 98.29 | 99.14 |
| Ramachandran allowed (%) | 2.84 | 3.23 | 6.49 | 1.71 | 0.86 |
| Ramachandran outliers (%) | 0.24 | 0.00 | 0.00 | 0.00 | 0.00 |
| Average B-factor | 98 | 51 | 81 | 57 | 52 |
| macromolecules | 98 | 51 | 81 | 57 | 52 |
| Solvent | n/a | 42 | n/a | 49 | 48 |

Statistics for the highest-resolution shell are shown in parentheses.
